# Defining the genetic landscape of acid and oxidative stress tolerance in *Streptococcus mutans* by pooled CRISPR interference screening

**DOI:** 10.64898/2026.08.04.742890

**Authors:** Yaqi Chi, Yuxing Chen, Chongyang Yuan, Liuchang Yang, Mingrui Zhang, Xiaolin Chen, Yiran Zhao, Ming Li, Xiaoyan Wang, Yongliang Li

## Abstract

Bacterial stress tolerance is a fundamental ecological adaptation that enables survival, competitive fitness, and persistent colonization under fluctuating environmental conditions. Dental caries remains a major global health burden. As an etiologic agent in caries, *Streptococcus mutans* (*S. mutans*) has the ability to adapt to sudden and substantial acid and oxidative stress. Nevertheless, the genome-wide genetic programs that support these stress tolerances remain incompletely defined. Herein, we established a xylose-inducible, genome-wide pooled CRISPR interference (CRISPRi) platform in *S. mutans* and performed parallel functional screens under acidic conditions (pH 5.0) and low-dose hydrogen peroxide (H□O□) stress. Using predefined screening thresholds, we identified 422 genes whose repression reduced fitness during acid challenge and 337 genes whose repression reduced fitness during H□O□ exposure. Functional and network analyses revealed that stress tolerance is strongly constrained by core physiological processes, including RNA (particularly transfer RNA) metabolism, macromolecule maintenance, and damage repair. Comparative analyses further indicated that growth-associated pathways displayed opposite trends between the two stresses, consistent with a stress-dependent allocation trade-off. These two complementary findings redefine the dual-stress adaptation paradigm of *S. mutans*: stress resistance is not determined merely by canonical stress signaling, but requires intact core physiological modules, and bacteria tune resource allocation dynamically to adapt to divergent stress microenvironments. Collectively, the present study provides a comparative, genome-scale functional map of acid- and oxidative-stress tolerance in a key oral pathogen and identifies genetic determinants for mechanistic studies and anti-caries interventions.

## Introduction

In natural and host-associated ecosystems, microorganisms do not grow in constant environments but continuously encounter multiple selective pressures arising from nutrient fluctuations, chemical stresses, and community-level competition^[1]^. For bacteria, stress tolerance is not an isolated defensive response, but an ecological adaptation strategy^[2]^. Therefore, deciphering the genetic basis of bacterial stress tolerance is conducive to understanding the stability of microbial communities, the mechanisms underlying the formation of dominant species, and the colonization and persistence of pathogenic bacteria within host ecological niches.

Dental caries is one of the most prevalent diseases worldwide, with bacterial dysbiosis recognized as a major contributor to its development^[3]^. Notably, *Streptococcus mutans* (*S. mutans*) is considered a key etiological agent due to its ability to adhere to tooth surfaces, form a robust biofilm matrix, ferment dietary carbohydrates into organic acids, and drive the transition from non-cariogenic to cariogenic biofilm states^[4, 5^]. To sustain this pathogenic lifestyle, *S. mutans* must repeatedly withstand two dominant selective pressures within the oral niche. First, it encounters oxidative stress generated by peroxigenic commensals in non-cariogenic biofilms^[6, 7^] and by host-derived reactive oxygen^[8]^. These oxidants damage DNA, proteins, and membrane lipids, thereby constraining bacterial survival and virulence-related functions^[9]^. Second, frequent sugar fermentation and limited buffering within mature biofilms cause recurrent and prolonged pH drops^[5]^. Consequently, *S. mutans* must possess strong tolerance to acid stress to survive.

Previous studies have largely attributed stress tolerance to specific stress-response pathways and key protective factors^[10]^, including F-ATPase-mediated H^+^ extrusion^[11, 12^], unsaturated fatty acid biosynthesis^[13, 14^], and the agmatine deiminase system (AgDS)-mediated pH homeostasis^[15, 16^]. Meanwhile, oxidative stress resistance is supported by enzymatic antioxidants (such as superoxide dismutase and catalase)^[17]^ and non-enzymatic redox buffers, including glutathione^[18]^, to scavenge ROS and mitigate oxidative damage^[8]^. Of note, our previous work showed that core housekeeping proteins, such as FtsZ, can maintain their protein activity under acidic conditions, thereby promoting the survival of *S. mutans* at low pH^[19]^. Moreover, accumulating evidence across a wide range of bacterial species indicates that stress adaptation can involve multiple basic physiological processes. Thus, under long-term stressors such as acidification, the competitive fitness of *S. mutans* may rely more heavily on the maintenance of homeostasis within core physiological systems than on the exclusive contribution of a limited set of canonical stress pathways.

Numerous studies have previously applied diverse omics approaches, including transcriptome sequencing^[20]^, proteomic profiling^[21]^, and metabolomics^[22]^, to profile stress-responsive pathways and candidate genes linked to these tolerance phenotypes. Nonetheless, such datasets are largely correlative, making it difficult to distinguish causal drivers from downstream consequences and to pinpoint genes that directly determine tolerance. Recently, CRISPR interference (CRISPRi)-based functional screens have enabled genome-scale interrogation of gene–phenotype associations in bacteria, providing quantitative landscapes of genetic determinants of stress tolerance^[23]^ and antibiotic resistance^[24]^. However, existing CRISPRi resources in *S. mutans* have largely relied on arrayed libraries targeting a limited set of genes (approximately 250 genes)^[25]^, restricting comprehensive genome-wide analyses.

To address this gap, the present study established a xylose-inducible, genome-wide pooled CRISPRi platform in *S. mutans* and used it to systematically interrogate the contributions of genes to acid tolerance and hydrogen peroxide (H□O□) tolerance. Furthermore, stress-specific screens, functional enrichment, and network-based analyses were combined to define stress-specific genetic programs. Importantly, our framework enables a comparative, genome-scale view of stress adaptation in *S. mutans* and serves as a resource for prioritizing genetic nodes for mechanistic follow-up.

## Materials and Methods

### 2.1 Bacterial strains, plasmids and growth conditions

*Streptococcus mutans* UA159 was obtained from the American Type Culture Collection (ATCC). Strains and plasmids used in this study are listed in Table S1. Strains were stored in 20% (*v/v*) glycerol at −80□°C. For routine cultivation, strains were streaked on brain heart infusion (BHI) agar and incubated at 37□°C for 24–48□h in 5% CO□. Single colonies were inoculated into CDY broth (chemically defined medium [CDM] supplemented with 0.3% yeast extract; Oxoid, UK) for subsequent experiments. For biofilm assays, CDYS (CDY supplemented with 1% (*w/v*) sucrose) was used.

### 2.2 Construction of the *S. mutans* pooled CRISPRi library

Here, sgRNAs were designed using the CRISPRi-seq scripts developed by Bakker *et al*. (https://github.com/veeninglab/CRISPRi-seq) based on the *S. mutans* UA159 genome (NCBI RefSeq assembly GCF_000007465.2). The PAM was set to NGG. For each gene, one sgRNA was selected, and oligonucleotides were synthesized with BbsI-compatible overhangs (forward: ATGT; reverse: AAAC). In total, 1,919 sgRNA pairs were synthesized (Beijing Ruibo Xingke Biotechnology Co., Ltd.).

A pooled sgRNA library was constructed as described previously^[26]^. Briefly, single-stranded oligonucleotides were annealed to generate double-stranded inserts and mixed at equimolar ratios. The pooled inserts were phosphorylated using T4 polynucleotide kinase (NEB M0525S) and ligated into the BbsI-digested pYL02 vector using T4 DNA ligase. The pYL02 is a suicide plasmid for *S. mutans* (sequence provided in the Supplementary Materials) that contains an ampicillin resistance marker and an *E. coli* replication origin for propagation in *E. coli*, and an erythromycin resistance cassette together with an sgRNA expression cassette. sgRNA expression is driven by the P3 promoter, and erythromycin resistance is driven by the P23 promoter. The sgRNA and erythromycin resistance cassettes were then flanked by two homology arms to enable chromosomal integration via homologous recombination.

Ligation products were electroporated into *E. coli* Stbl3 competent cells (Transgene) and plated on LB agar containing ampicillin (100□µg□mL□¹). After overnight incubation, approximately 2.1□×□10□ colonies were collected, resuspended in LB medium and thoroughly mixed. An aliquot (5□mL) was mixed with an equal volume of 50% glycerol (v/v) and stored at −80□°C. The remaining suspension was used for plasmid extraction (Plasmid Maxi Kit, Qiagen).

To assess sgRNA representation in the plasmid pool, the sgRNA region was PCR-amplified using Phanta high-fidelity DNA polymerase (Vazyme, Beijing). Amplicons were purified and quantified (Qubit dsDNA HS Assay Kit) and sequenced on an Illumina MiniSeq. The sgRNA abundance was quantified from processed reads using MAGeCK.

To generate a xylose-inducible CRISPRi host strain, the endogenous Cas9 was deleted by in-frame deletion. An exogenous dspCas9_FLAG gene under control of a TC-Xyl-inducible cassette was integrated into the phnA–mtlA locus by overlapping PCR and homologous recombination as described previously^[27, 28^]. The pooled sgRNA plasmid library was introduced into this strain via CSP-mediated transformation^[27]^, and the sgRNA cassette was integrated at the phnA–mtlA region. Transformants were selected on BHI agar containing erythromycin (12□µg□mL□¹) (Figure 1). The *ftsW*- and *pbp2x*-targeting CRISPRi strains were constructed using the same workflow, except that individual sgRNAs targeting *ftsW* or *pbp2x* replaced the pooled library.

**Figure 1.**
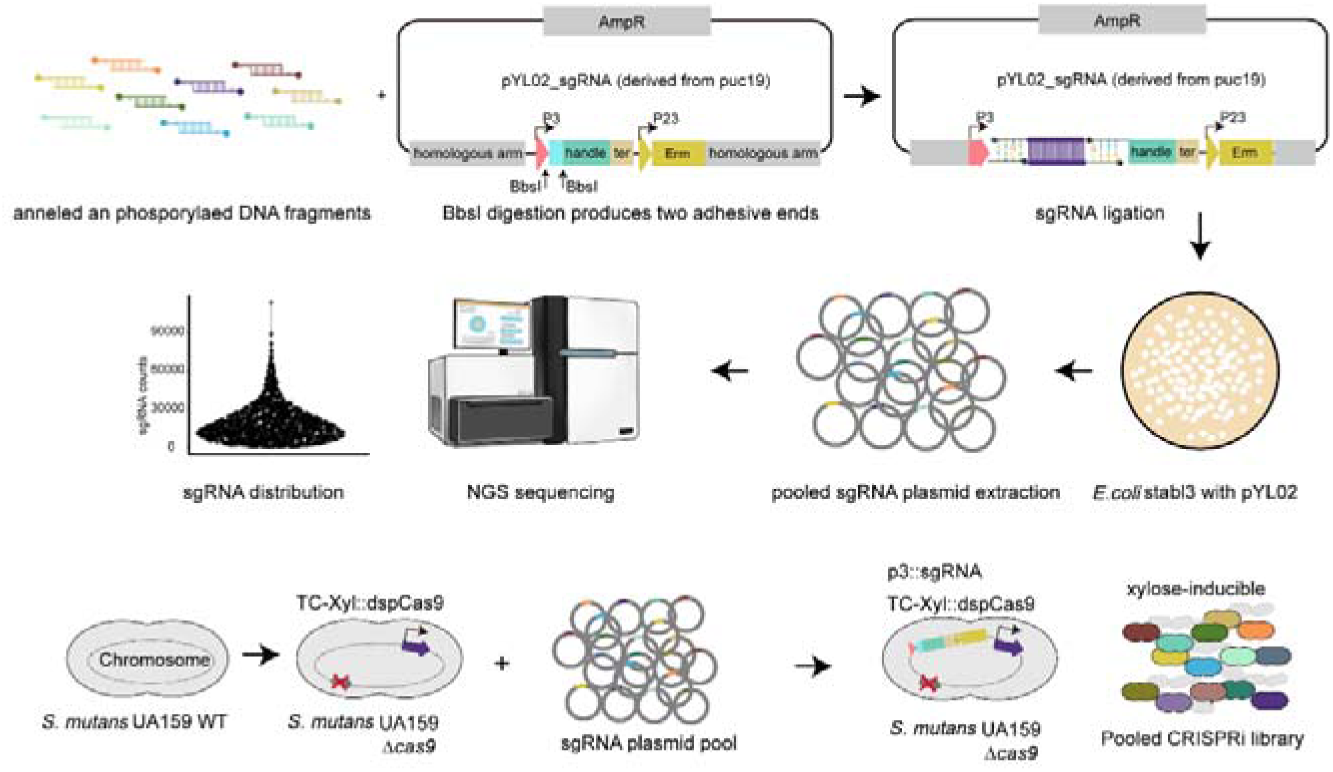
Construction of a genome-wide pooled CRISPRi library in *S. mutans*. Schematic overview of library design and assembly. We designed 1,919 sgRNAs targeting the S. mutans genome. The pYL02 vector was digested and ligated with the phosphorylated sgRNA oligonucleotide pool, and the ligation products were electroporated into *E. coli* Stbl3 to amplify the plasmid library. The pooled sgRNA plasmids were recovered from transformants, and next-generation sequencing showed an approximately normal distribution of sgRNA abundances. In the recipient S. mutans strain, the endogenous cas9 was deleted by in-frame deletion, and a TC-Xyl-inducible dspCas9 cassette was integrated into the chromosome. The pooled sgRNA plasmid library was then introduced into this strain via CSP-mediated transformation.

### 2.3 CRISPRi-seq

#### 2.3.1 Sample preparation

Genome-wide screens were performed using the pooled sgRNA library. The library was resuscitated in CDY broth (1:20 dilution), cultured to mid-log phase, and subsequently inoculated into four experimental conditions: (1) CDY, pH 7.4; (2) CDY supplemented with 1% (*w/v*) xylose, pH 7.4; (3) CDY supplemented with 1% (*w/v*) xylose, pH 5.0; and (4) CDY supplemented with 1% (*w/v*) xylose and 0.003% (*v/v*) H□O□, pH 7.4 (Figure S1).

To identify genes associated with growth under induction, cultures in conditions (1) and (2) were serially passaged three times, with each passage initiated from mid-log phase cultures, and essentiality under induction was inferred by comparing (2) versus (1). Acid tolerance genes were identified by comparing condition (3) versus condition (2) using the serially passaged samples (Figure 3A). Oxidative stress tolerance-associated genes were identified by culturing the library under conditions (2) and (4) for a single passage to mid-log phase without serial passaging, followed by comparison of condition (4) versus condition (2) (Figure 4A). Total genomic DNA was extracted using the TIANamp Bacteria DNA Kit (Tiangen, China). DNA concentration and purity were assessed using a NanoDrop ND-1000 spectrophotometer (Thermo Fisher Scientific).

**Figure 2.**
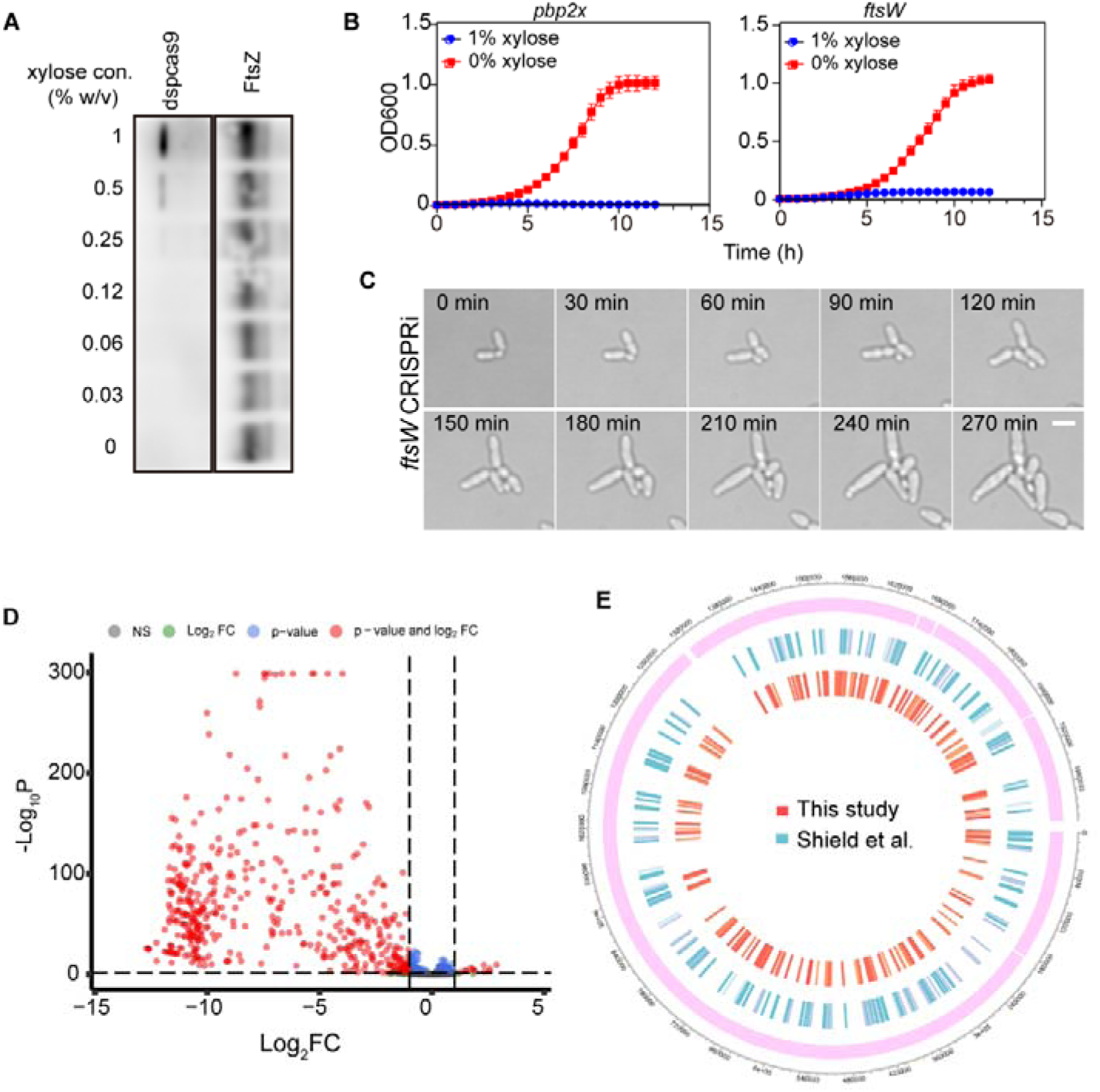
Validation of the CRISPRi library. (A) Western blot analysis of dspCas9 expression in *ftsW* CRISPRi strains under increasing xylose concentrations (0–1% w/v). FtsZ was used as a loading control. DspCas9 expression was induced in a dose-dependent manner, with no detectable expression in the absence of xylose. (B) Volcano plot of essential genes. (C) Growth curve of the *ftsW* and *pbp2x* CRISPRi strain under 1% xylose-induced or uninduced conditions. (D) Cell division images of the *ftsW* CRISPRi strain under non-inducing conditions. (E) The genomic distribution of essential genes identified in this study was visualized and compared with those previously reported by Shield *et al*. via Tn-seq using a Circos plot. The outermost ring represents the reference genome, with inner tracks displaying essential genes identified in this study (red bars) and those from Shield *et al*. (blue bars) at their corresponding genomic positions.

**Figure 3.**
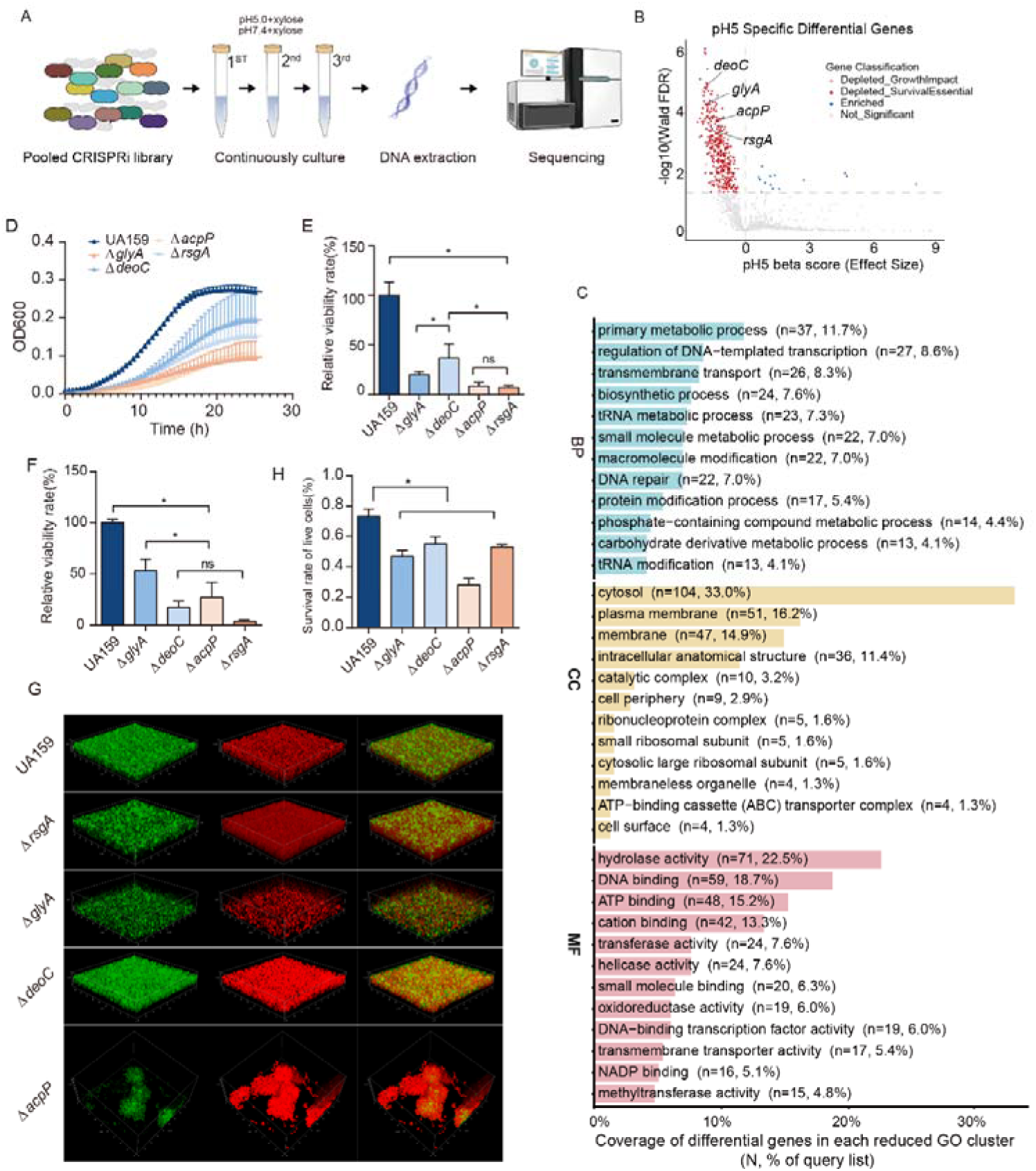
Genome-scale CRISPRi screening reveals essential genes and acid-tolerant genes. (A) Schematic overview for CRISPRi screening in *S. mutans*. NGS was used to determine the changes in sgRNA abundance. (B) Volcano plot of acid-tolerant genes. NGS sgRNA abundance comparison of CRISPRi library exposed to acid condition (pH = 5.0, 1% xylose) or to the control condition (pH = 7.4, 1% xylose) (*n*□=□3 for each condition). (C) Gene functions were categorized using annotations from the Gene Ontology (GO) database. (D) Growth curves of UA159, Δ*glyA*, Δ*deoC*, Δ*acpP*, and Δ*rsgA* strains under acidic conditions (pH = 5.0). (E) Relative viability rate of UA159, Δ*glyA*, Δ*deoC*, Δ*acpP*, and Δ*rsgA* strains in a direct acid kill assay. (F) Relative viability rate of UA159, Δ*glyA*, Δ*deoC*, Δ*acpP*, and Δ*rsgA* strains in acid-tolerant response assay. (D) Survival rates of live cells of WT and mutants. (G) CLSM images of 48-h biofilms formed by the UA159, Δ*glyA,* Δ*deoC,* Δ*acpP, and* Δ*rsgA* strains. The green fluorescence indicates live cells, while the red fluorescence indicates dead cells. Images were examined at 60 × objective magnification. Scale bar: 10 µm (*, *P*□<□0.05; ns, not significant).

**Figure 4.**
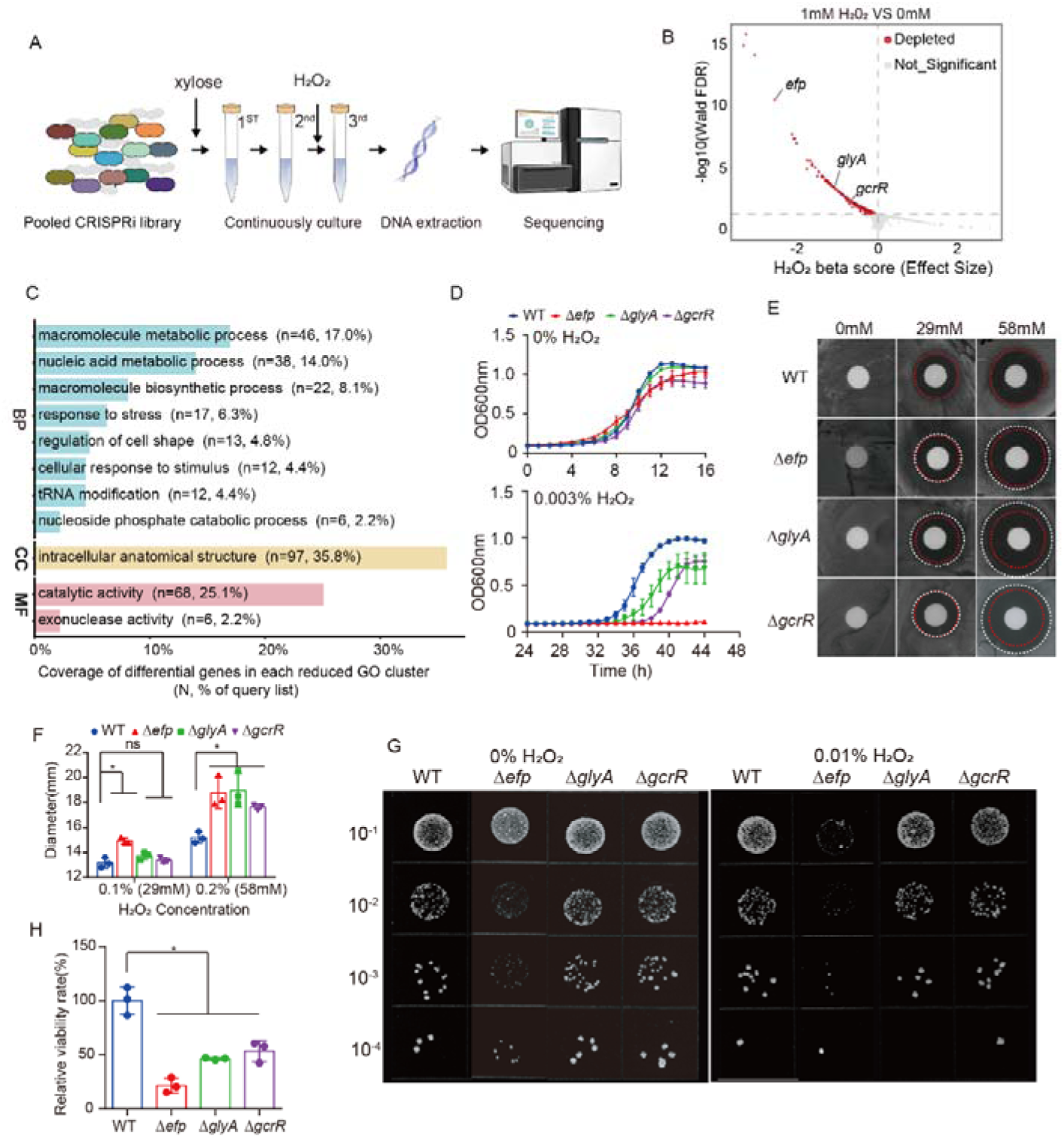
Genome-scale CRISPRi screening reveals oxidative stress tolerance genes. (A) Schematic overview for CRISPRi screening in *S. mutans*. NGS was used to determine the changes in sgRNA abundance. (B) Volcano plot of oxidative stress tolerance genes. NGS sgRNA abundance comparison of CRISPRi library exposed to oxidative stress condition (0.003% H_2_O_2_, 1% xylose) or to the control condition (0% H_2_O_2_, 1% xylose) (*n*□=□3 for each condition). (C) Gene functions were categorized using annotations from the Gene Ontology (GO) database. (D) Growth curves of UA159, Δ*glyA*, Δ*efp,* and Δ*gcrR* strains under oxidative stress (0.003% H_2_O_2_). (E, F) Representative images (E) and diameter measurement (F) of inhibition zones formed by UA159, Δ*glyA*, Δ*efp*, and Δ*gcrR* mutant strains in response to 29□mM and 58□mM H_2_O_2_ (*, *P*□<□0.05; ns, not significant). (G, H) Representative images (G) and (H) relative viability of *S. mutans* UA159, Δ*glyA,* Δ*efp, and* Δ*gcrR* strains following treatment with 0.01% H_2_O_2_ for 30 min. Relative survival was determined by CFU counting and normalized to UA159.

#### 2.3.2 sgRNA sequencing and data analysis

sgRNA libraries were prepared from extracted DNA using a two-step PCR amplification procedure as described previously^[29]^. In the first PCR step, the sgRNA cassette was amplified from the library. In the second PCR step, an 8-bp sample barcode and a stagger sequence were added to enable multiplexing and increase sequence complexity^[29]^. Primers and cycling conditions are provided in the Supplementary Materials. Second-round PCR products were purified, pooled, diluted, spiked with 10% PhiX, and sequenced on an Illumina NovaSeq 6000 platform.

Low-quality reads were removed using fastp v0.20.0 as described previously^[30]^. Read counts were normalized to counts per million (CPM). Gene-level fitness effects were estimated using the MAGeCK-MLE model, which reports effect sizes (beta) and statistical significance (wald-fdr) based on a Wald test with Benjamini–Hochberg correction. Genes were considered significantly depleted at wald-fdr < 0.05. To identify stress-specific candidates, genes with significant negative fitness under induction were excluded from the pH- and H□O□-specific hit lists. For interpretation, the induced-to-uninduced read ratio was calculated. Ratios of 0.01–0.1 indicated compromised fitness, whereas ratios <0.01 were considered consistent with essentiality, as described previously^[31]^. Functional categorization was performed using Gene Ontology enrichment and annotation resources^[32, 33^]. GSEA was performed using ranked gene lists based on MAGeCK-MLE beta values, including pH 5.0 and H□O□ datasets and combined scores for same-direction and opposite-direction trends. GSEA was implemented using fgseaMultilevel, with *P* values adjusted by the Benjamini–Hochberg method; FDR ≤ 0.1 was used as the significance threshold^[34]^. Protein–protein interaction networks were obtained from STRING and analyzed in Cytoscape v3.10.3.

### 2.4 Construction of *S. mutans* mutants

Null mutants (Δ*glyA*, Δ*deoC*, Δ*acpP*, Δ*rsgA*, Δ*efp*, Δ*pbp2b*, and Δ*gcrR*) were generated in *S. mutans* UA159 using the IFDC2 cassette by overlapping PCR and homologous recombination, as described previously^[27, 28^]. Primers are listed in Table S2.

### 2.5 Western blotting

The *ftsW* CRISPRi strain was grown to mid-log phase in CDY medium supplemented with xylose (0, 0.03, 0.06, 0.12, 0.25, 0.5, and 1% (*w/v*)). Total protein was extracted, separated by SDS–PAGE, and transferred onto PVDF membranes. Immunoblotting was performed using anti-FLAG (30504ES50, YEASEN, China) and anti-FtsZ (custom-made) primary antibodies, followed by appropriate secondary antibodies. Signals were detected by chemiluminescence (GE AI600).

### 2.6 Live-cell imaging of the *ftsW* CRISPRi strain

The *ftsW* CRISPRi strain was cultured overnight in CDY medium at 37□°C, diluted 1:100 into CDY medium containing 1% (*w/v*) xylose, and grown to OD□□□□=□0.4–0.5. Cultures were rediluted 1:100 in the same medium and grown to OD□□□□=□0.1–0.2. Agarose pads were prepared with 1.5% low-melting-point agarose in CDY medium containing 1% xylose.

Cells were imaged using an N-STORM system (Nikon, Tokyo, Japan) equipped with a 100× oil TIRF objective (Nikon Plan Apo, 1.49 NA), an Andor-897 EMCCD camera (Andor, Belfast, Northern Ireland), and 1.5× magnification optics. Bright-field images were acquired every 30□min for 270□min using NIS-Elements AR. Raw .nd2 files were converted to .tiff and processed in ImageJ (NIH, Bethesda, MD, USA).

### 2.7 Growth curve assay

Growth of UA159 wild type, *ftsW* CRISPRi, *pbp2x* CRISPRi, Δ*glyA*, Δ*deoC*, Δ*acpP*, Δ*rsgA*, Δ*efp*, and Δ*gcrR* was measured using a microplate reader (SpectraMax 190, Molecular Devices). Overnight cultures were diluted 1:100 in CDY medium, with uninoculated medium serving as the blank. OD□□□ was recorded every 30□min for 24–48□h after 5□s of shaking. Growth curves were generated from OD□□□ values.

### 2.8 Acid tolerance assays

Acid tolerance was assessed using a modified published protocol^[35]^. Log-phase cultures (OD□□□□=□0.5) were centrifuged (4,000 × *g*, 10□min, 4□°C) and washed once with 0.1□M glycine buffer (pH 7.0). Baseline CFUs were determined by dilution in PBS, plating on BHI agar, and incubation at 37□°C with 5% CO□ for 48□h. For constitutive tolerance, cells were exposed to 0.1□M glycine–HCl (pH 2.8) for 10□min. For the acid tolerance response (ATR), cells were incubated in CDY at pH 5.0 for 2□h before exposure to 0.1□M glycine–HCl (pH 2.8) for 10□min. Treated cells were diluted, plated, and incubated as above. Relative viability was calculated as follows:

Relative viability (%) = [(CFU_pH2.8 mutant / CFU_pH2.8 WT) / (CFU_control mutant 266 / CFU_control WT)] × 100.

### 2.9 Biofilm formation and confocal laser scanning microscopy

Log-phase cultures (OD□□□□=□0.5) of wild-type and mutant strains were diluted 1:100 in CDYS and incubated anaerobically at 37□°C for 48□h. Biofilms were imaged by CLSM using a 60× oil-immersion objective (TCS-SP8 and MICA, Leica). Excitation wavelengths were 488□nm for SYTO 9 and 561□nm for propidium iodide. Three randomly selected fields were imaged per biofilm.

### 2.10 H□O□ killing and sensitivity assays

H□O□ tolerance was assessed using a previously published protocol^[36]^, with conditions matched to those used in the killing assay. Procedures were analogous to the acid tolerance workflow, except that cells were exposed to 0.01% (2.9□mM) H□O□ for 30□min. For sensitivity testing, log-phase cultures (OD□□□□=□0.5) were spread (100□µL) on CDY agar. Filter discs containing 0, 29□mM (0.1%) or 58□mM (0.2%) H□O□ were placed on plates, followed by incubation at 37□°C for 24□h. Inhibition zones were measured at three positions, and the inhibition zone–to–disc diameter ratio was calculated.

### 2.11 AlphaFold structure prediction

Protein structure prediction was performed using AlphaFold3 (https://alphafoldserver.com/) with default parameters. The top-ranked model (ranked_0) was used for downstream analysis. Protein–protein interaction interfaces and oligomeric states were assessed using the PISA server (http://www.ebi.ac.uk/pdbe/pisa/). Structures were visualized in PyMOL (https://pymol.org/).

### 2.12 Statistical analysis

Statistical analyses were performed in GraphPad Prism 7 (v7.00). Variance homogeneity was assessed using Levene’s test. One-way ANOVA followed by the Student–Newman–Keuls (SNK) post hoc test was used for multiple comparisons. All tests were two-sided with α = 0.05.

### 2.13 Data availability

sgRNA sequencing data have been deposited in the Sequence Read Archive under accession PRJNA1470443 (https://dataview.ncbi.nlm.nih.gov/object/PRJNA1470443?reviewer=fp0a2mtj58ojdcm265n229jgal). Other datasets are available from the corresponding author upon reasonable request.

## Results

### 3.1 Construction of the pooled CRISPR interference library in *S. mutans*

We designed sgRNAs for *S. mutans* UA159 using a pipeline adapted from a previous study^[26]^. Candidate spacers were enumerated at SpCas9 NGG PAM sites previously validated for use in *S. mutans*, and were filtered using sequence constraints, including removal of BbsI sites and homopolymer runs. Off-target potential was evaluated by genome-wide matching with cumulative mismatch thresholds across the spacer and a position-weighted penalty scheme. From 49,694 candidate sgRNAs, an optimized set of 1,919 sgRNAs (one per protein-coding gene) was selected for pooled library construction, exhibiting a nearly 100% relative repression activity (Figure S2).

To validate inducible CRISPRi function, we first confirmed that the expression of dspCas9 was dependent on the concentration of xylose. As xylose concentrations increased from 0.03 to 1%, dspCas9 protein levels showed a gradual and dose-dependent upregulation (Figure 2A). We then constructed two CRISPRi strains targeting the essential genes *ftsW* and *pbp2x* using the optimized sgRNAs. In the presence of xylose, repression of either target effectively inhibited bacterial growth, consistent with efficient gene knockdown (Figure 2B). Live-cell imaging further revealed that repression of *ftsW* caused marked cell elongation and impaired cell division (Figure 2C). Taken together, these results validate the feasibility and functional efficacy of our CRISPRi-based gene repression system in *S. mutans*.

For pooled library construction, the optimized 1,919 sgRNAs were cloned into the pYL02 vector. After cloning, 2.1 × 10^6^ *E. coli* colonies were collected to maintain ≥1,000-fold coverage. Sequencing of the pooled plasmid library detected 1,896 sgRNAs, with an approximately normal abundance distribution. The library was then transformed into a UA159-derived strain, and 2 × 10^4^ colonies were collected (≥100-fold coverage). Using the pooled platform under inducing conditions, 399 sgRNAs showed significant depletion (Figure 2D). Genes with >100-fold depletion were defined as lethal essential genes, while genes with 2–100-fold depletion were characterized as growth-supporting genes. Based on these thresholds, a total of 213 essential genes and 186 growth-supporting genes were identified. Next, our essential gene set was compared with the Tn-seq dataset reported by Shields *et al*.^[31]^, which annotated 260 genes as essential or possibly essential. Notably, substantial concordance was observed at the genomic locus level (Figure 2E), with 173 genes overlapping with previously reported essential or potentially essential genes, while 226 genes were not documented as essential in the prior Tn-seq study (Table S3). This comparison indicates broad agreement between pooled CRISPRi and transposon-based essentiality mapping, while also highlighting method- and condition-dependent differences.

### 3.2 Genome-scale identification of acid tolerance determinants

Next, the pooled library was leveraged to identify acid resistance-related genes by comparing the sgRNA abundance under neutral (pH 7.4, CDY medium +1% xylose) and acidic conditions (pH 5.0, CDY medium +1% xylose) (Figure 3A). A total of 422 specifically depleted genes were identified under pH 5.0 (acidic stress) conditions (Table 1; Figure 3B). To enrich for acid-specific contributors, genes depleted under the inducing condition, regardless of pH, were excluded.

**Table 1.**
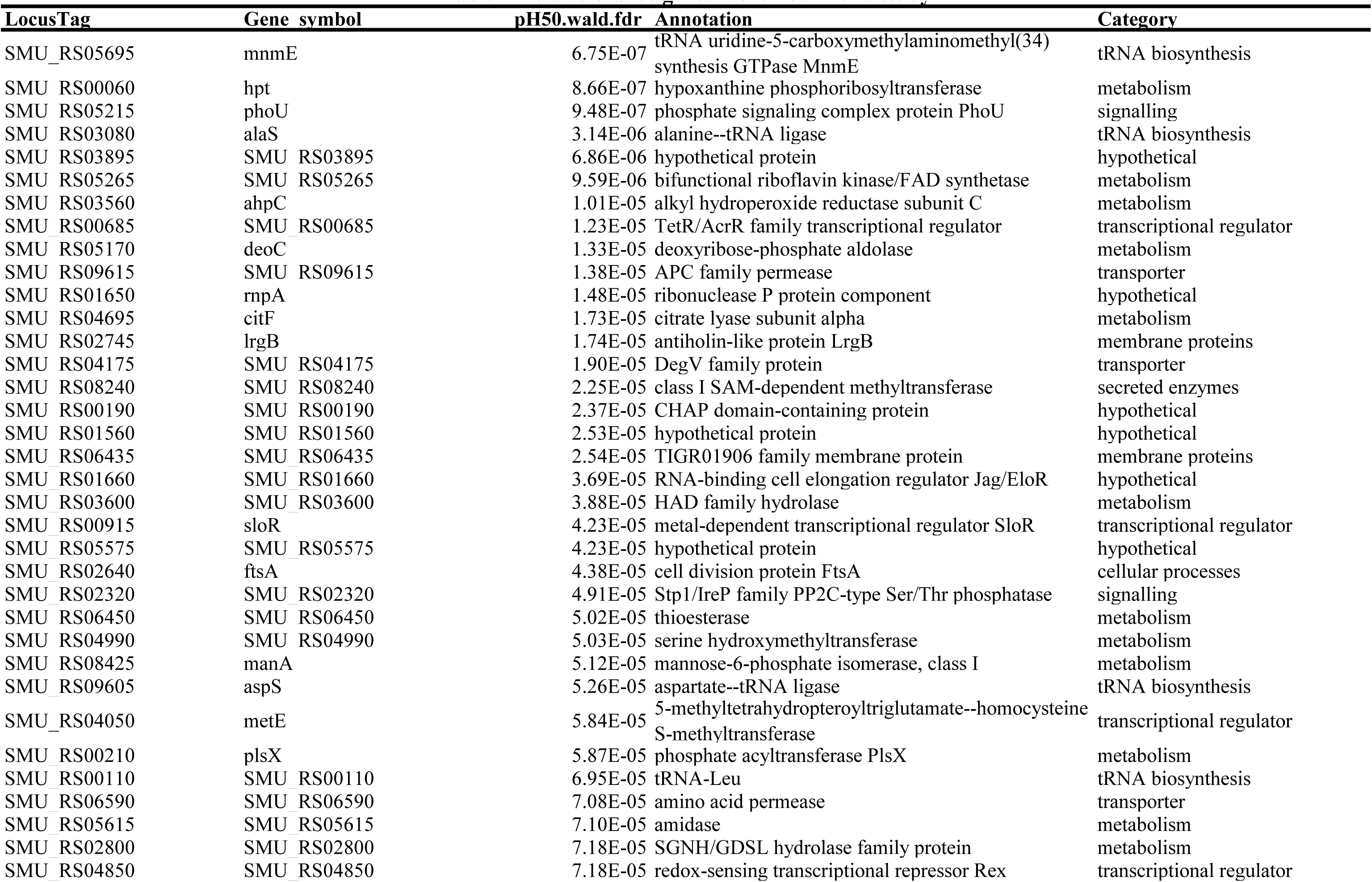
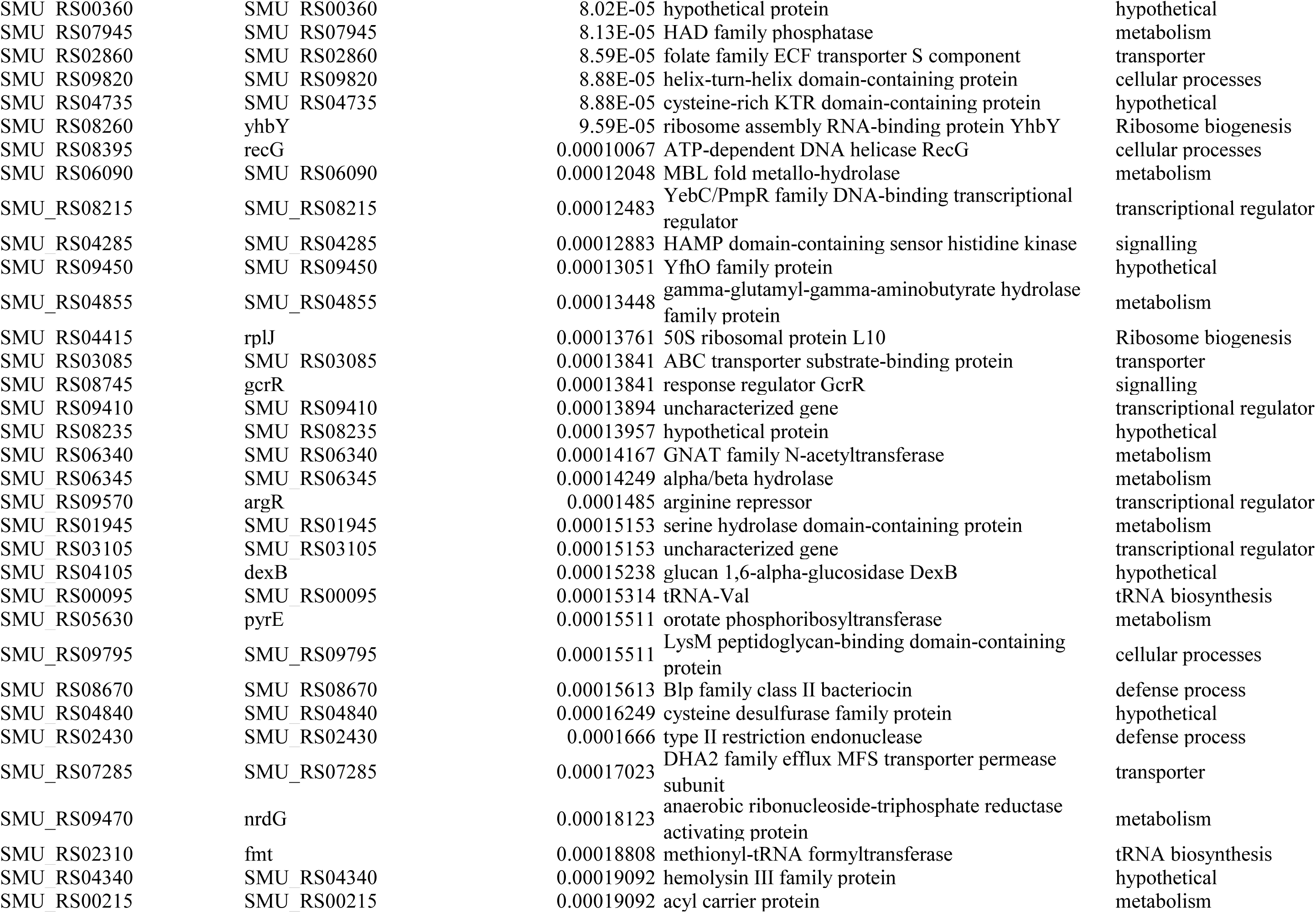
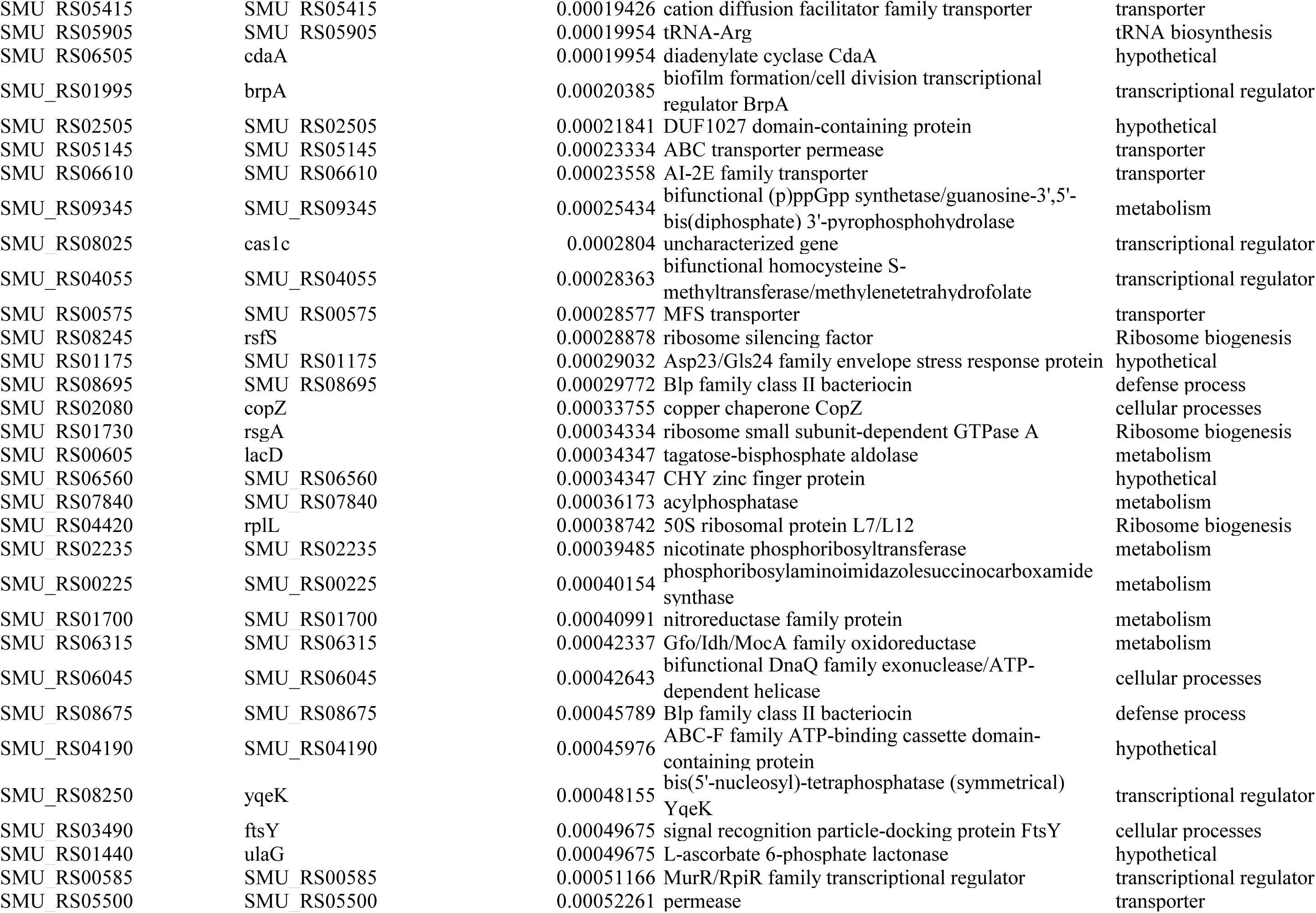
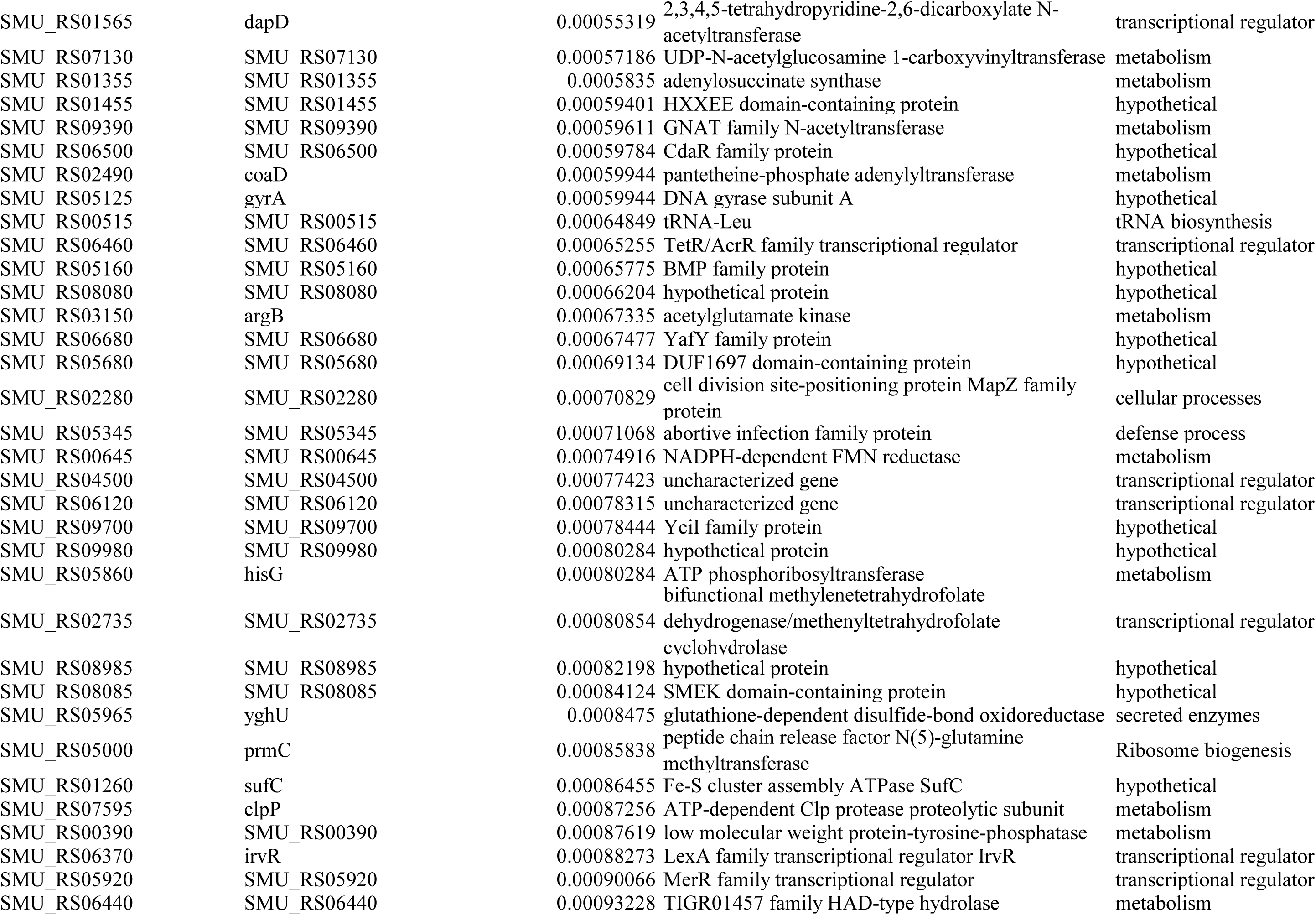
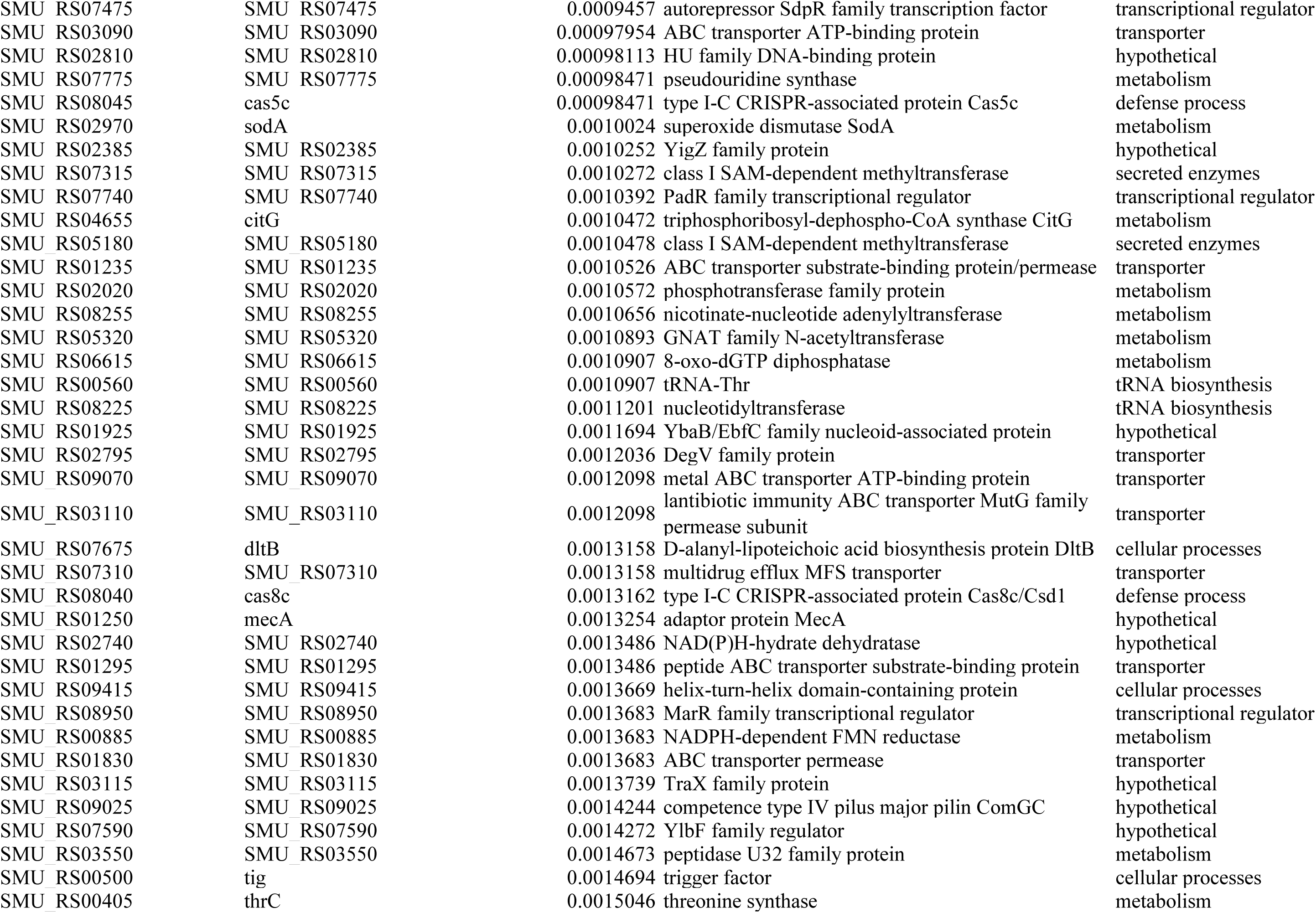
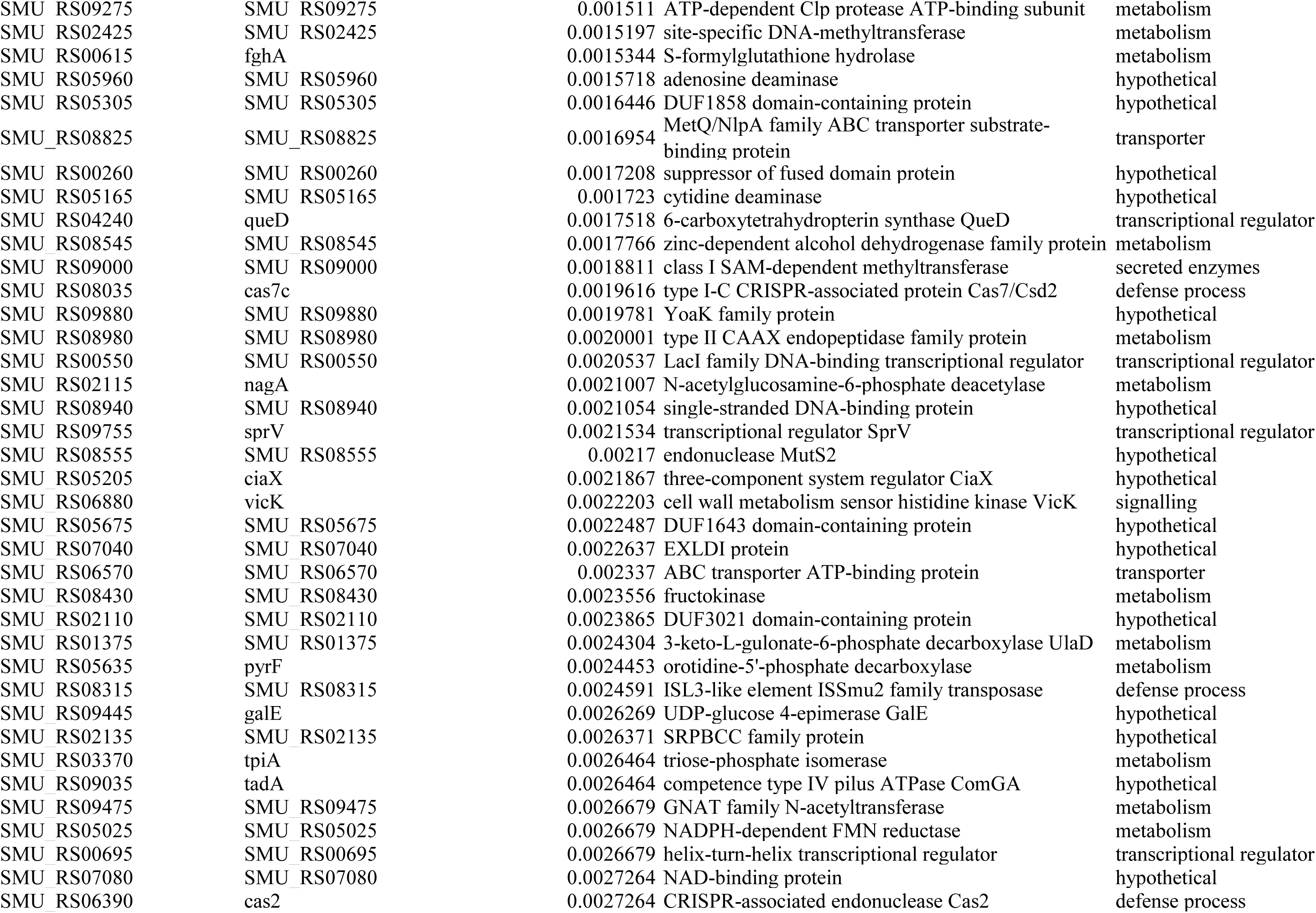
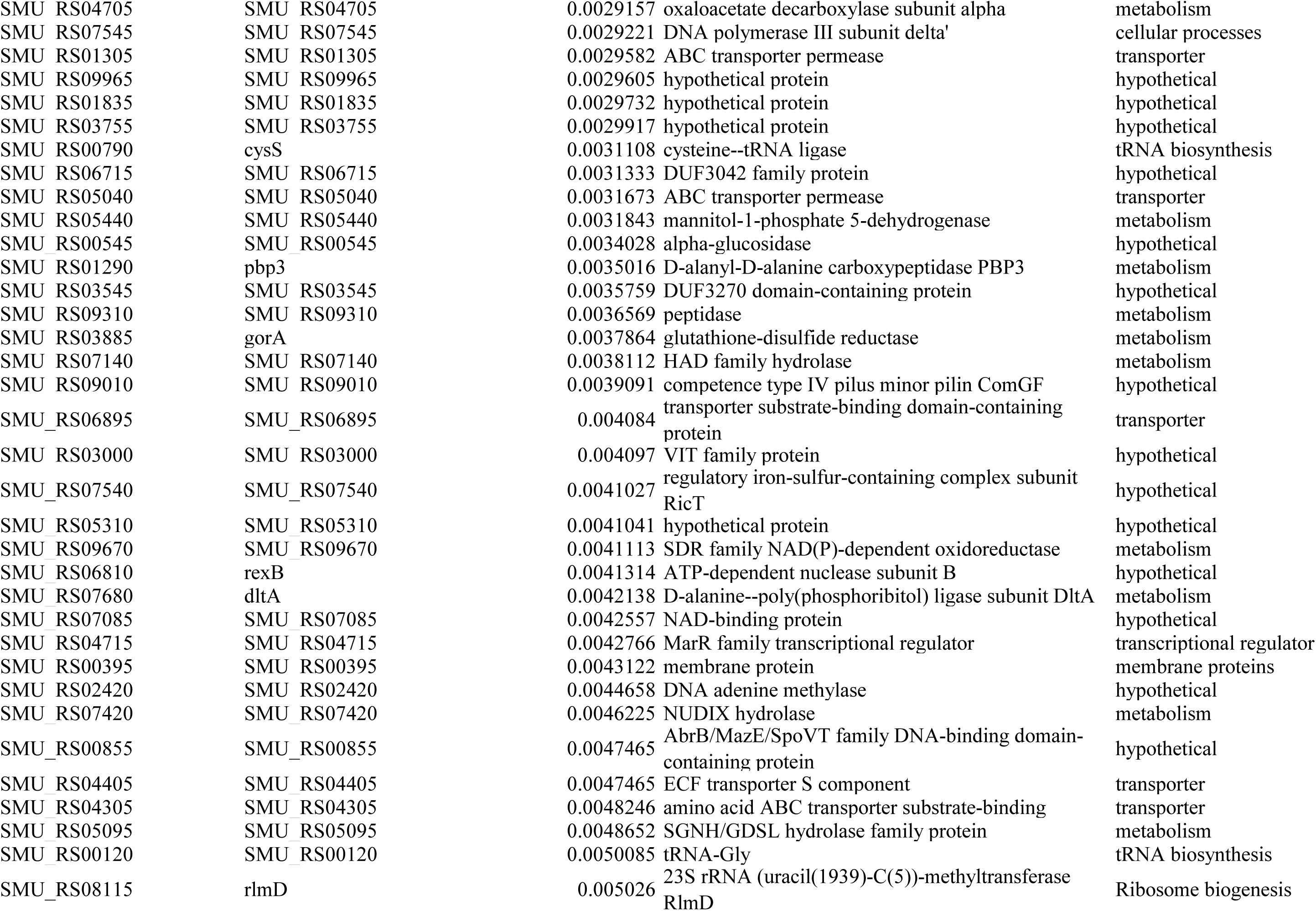
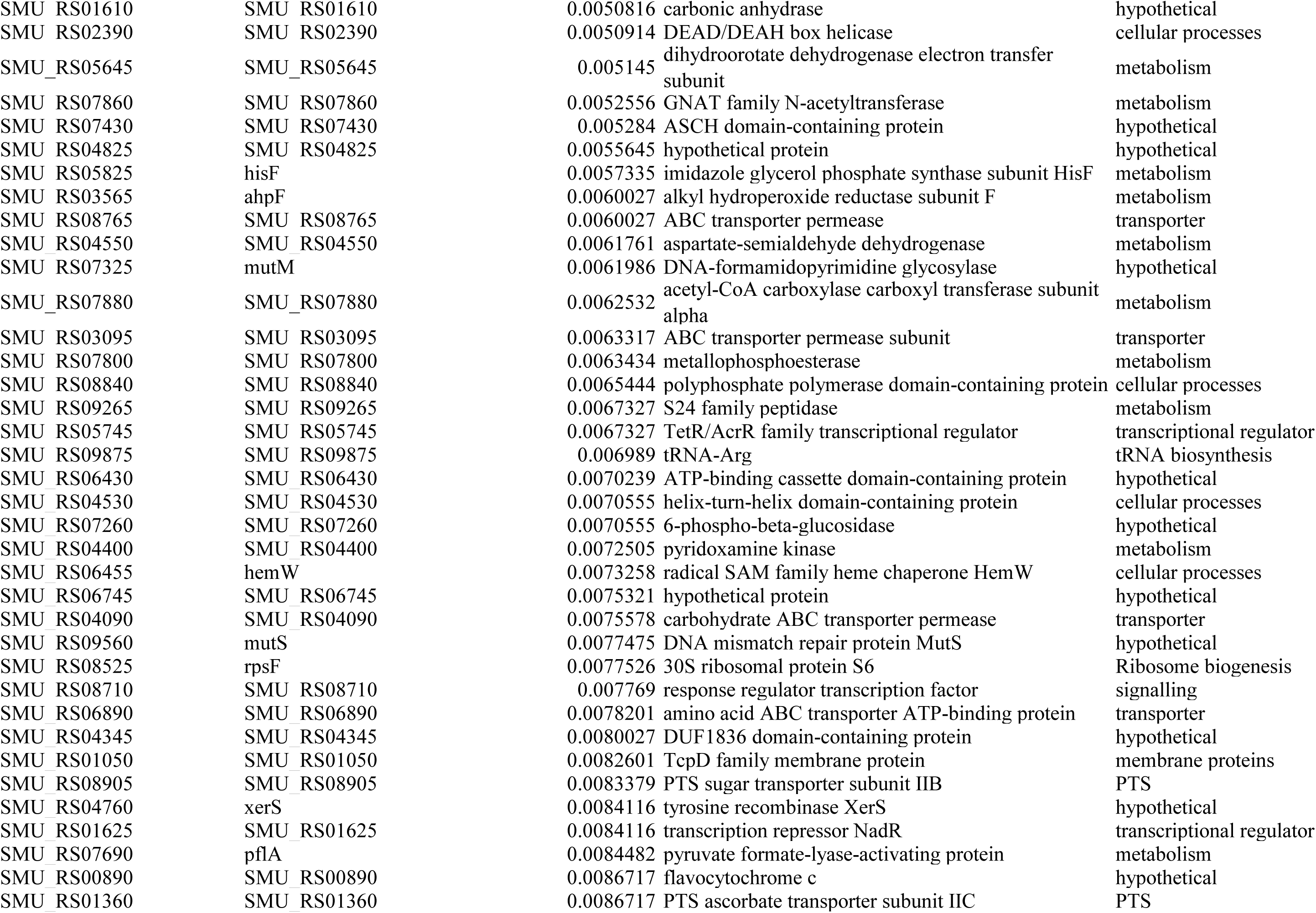
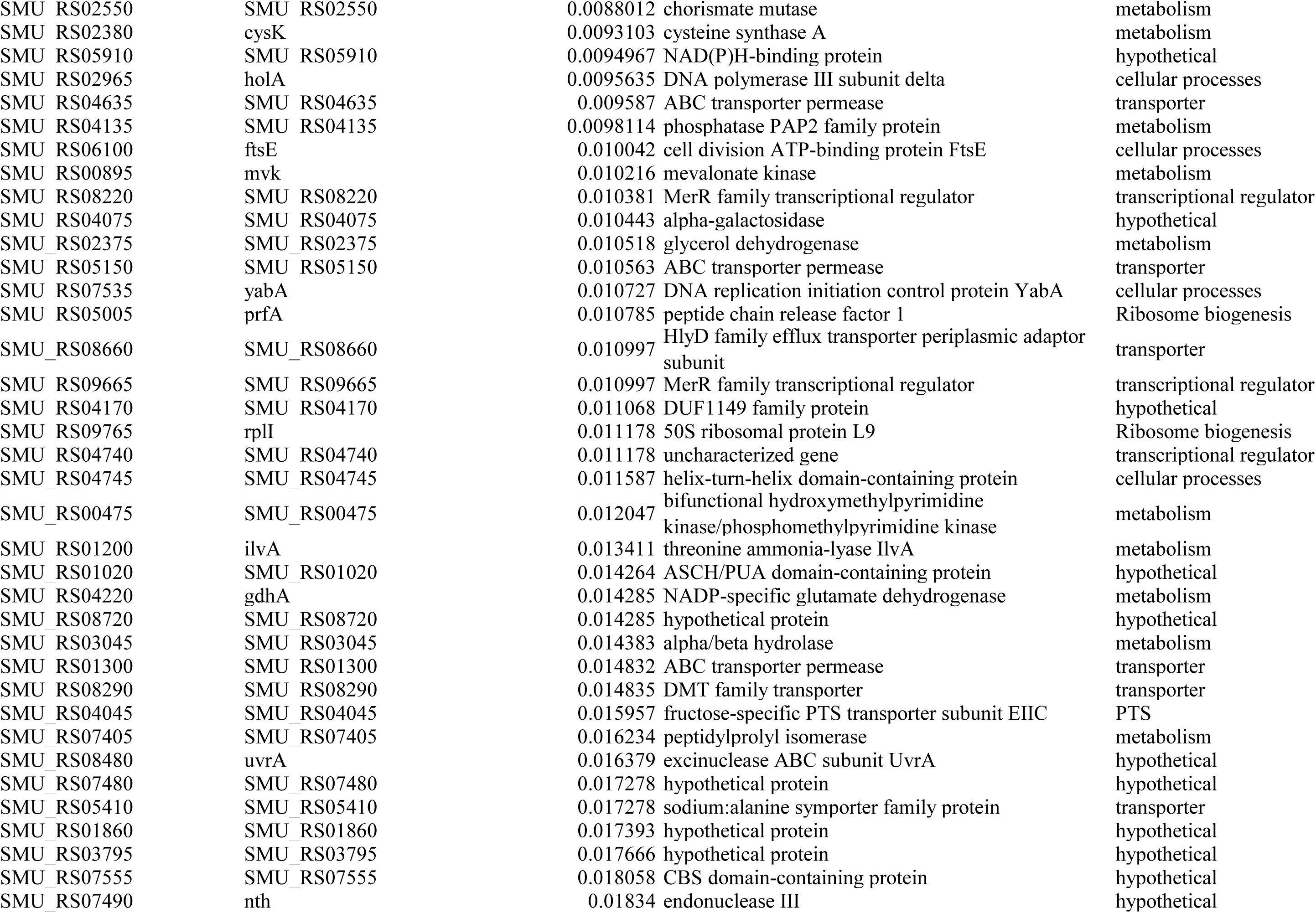
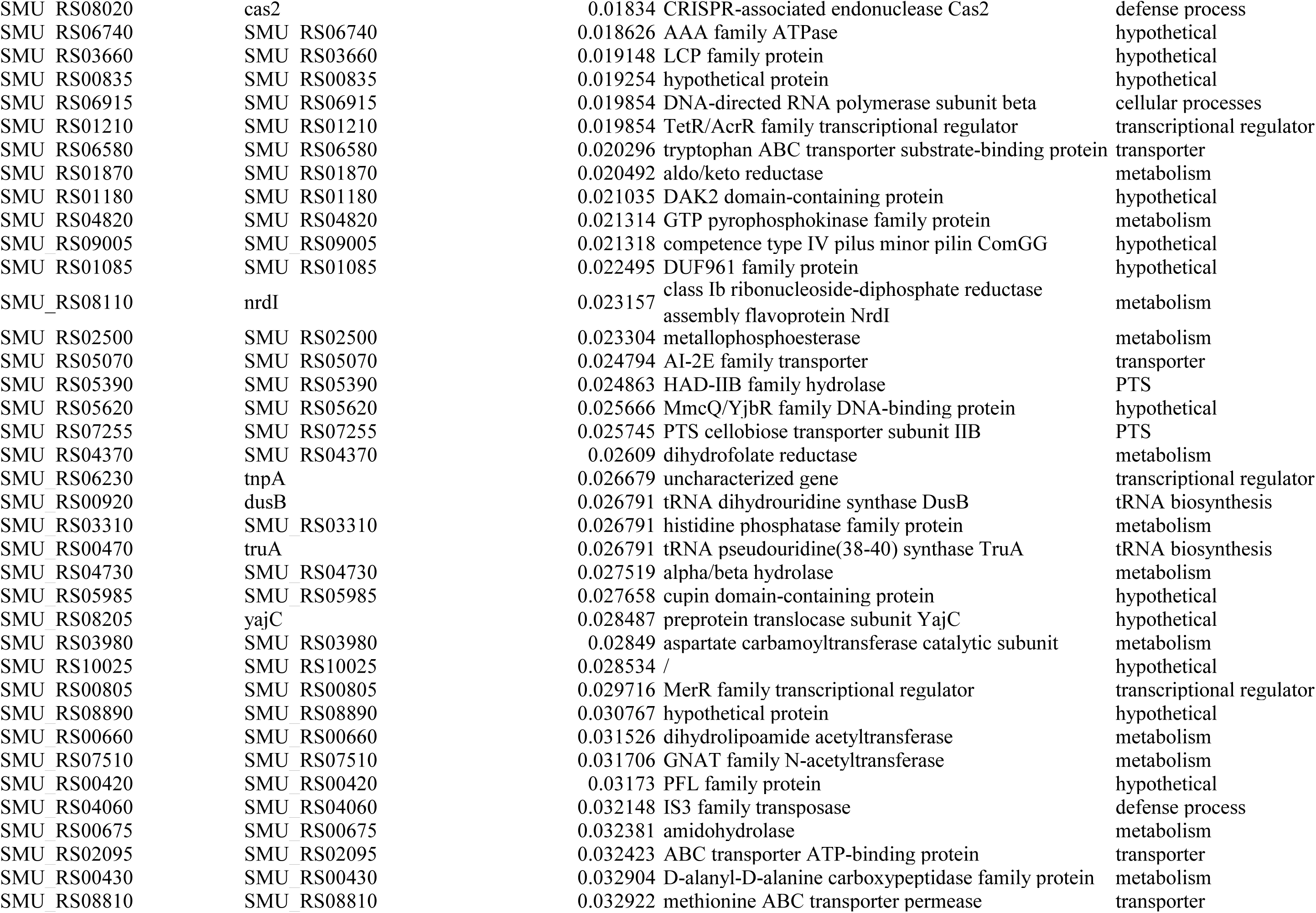
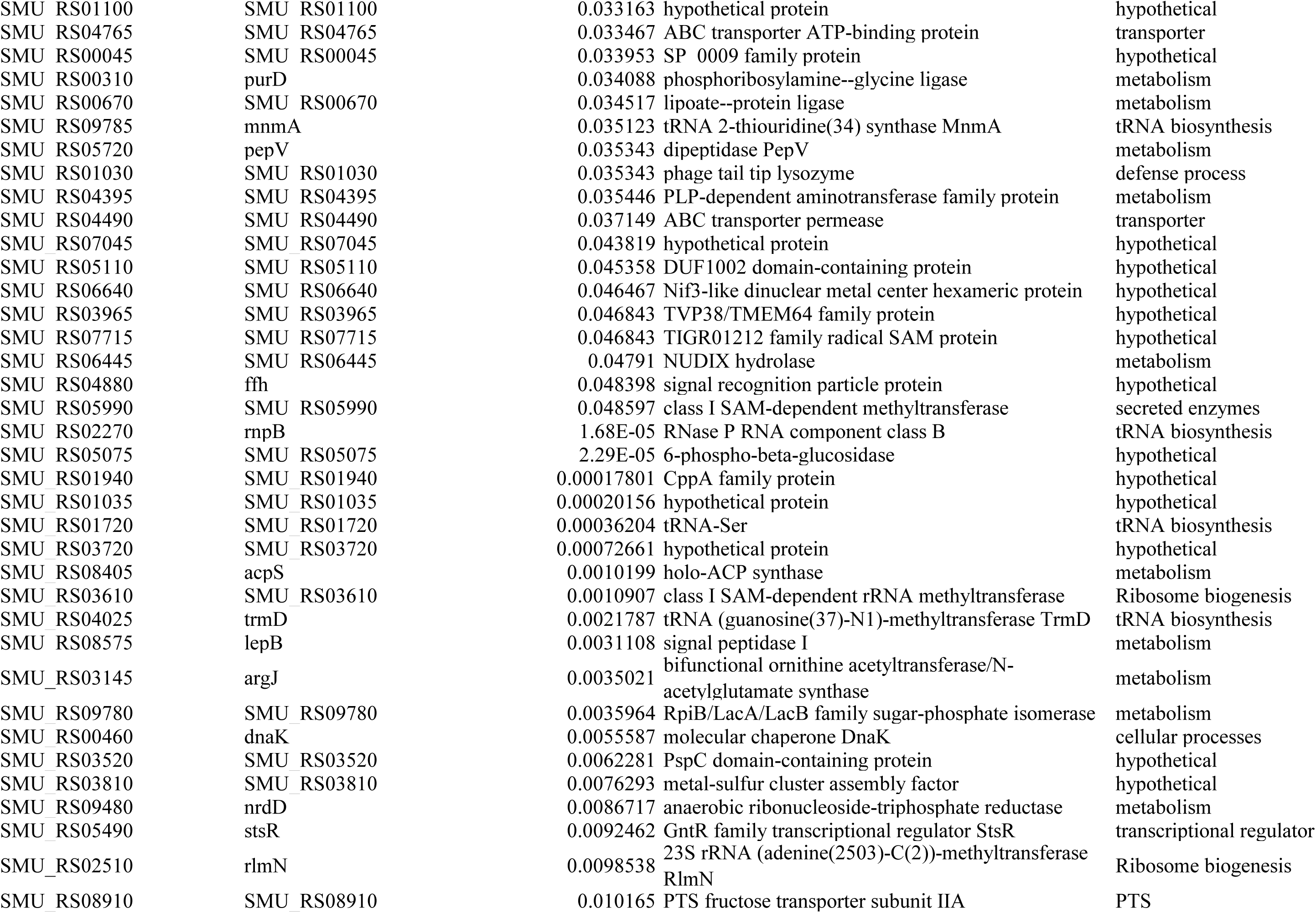
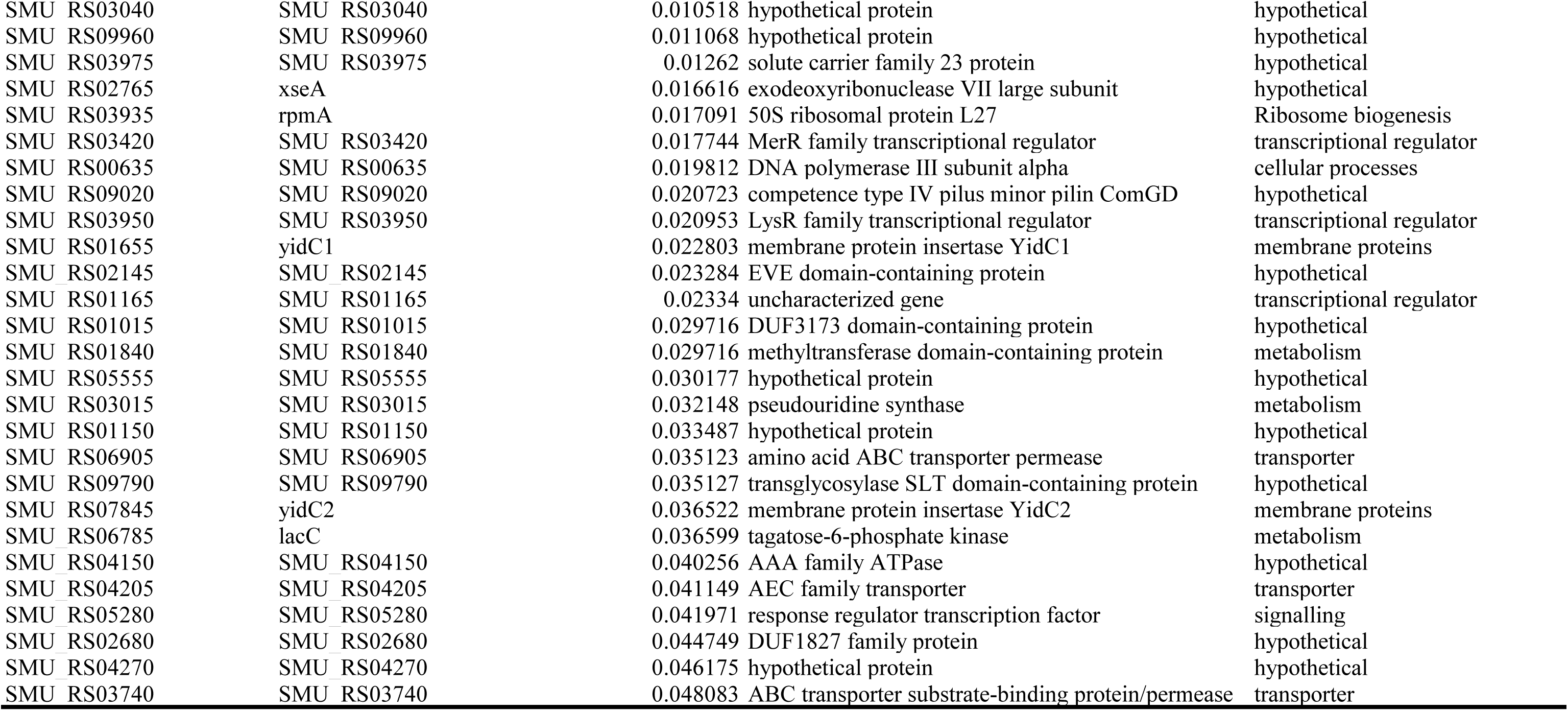
Acid-tolerance genes identified in this study.

Functional annotation revealed a broad requirement for cellular maintenance during acid challenge (Figure 3C). At the BP level, primary metabolic process (11.7%), transmembrane transport (8.3%), and DNA repair (7.0%) were prominent. Notably, tRNA metabolic/modification processes (7.3%/4.1%) and macromolecule modification (7.0%) were also enriched, suggesting a role for translational control and RNA/protein handling in acid adaptation. CC enrichment highlighted the cytosol (33.0%) and plasma membrane (16.2%), with transporter complexes further supporting a central role for membrane-associated functions. At the MF level, hydrolase activity (22.5%), DNA binding (18.7%), and ATP binding (15.2%) were dominant, with concurrent enrichment of cation binding (13.3%), consistent with coupling between energy metabolism and ion homeostasis.

To further validate the sequencing results, deletion mutants were generated for selected candidate genes (*glyA*, *deoC*, *acpP*, and *rsgA*) and their acid tolerance was assessed relative to the wild-type strain. Under acidic growth conditions (pH 5.0), mutants showed impaired growth, characterized by delayed entry into exponential phase and reduced final cell yields (Figure 3D). We then quantified both constitutive acid tolerance (Figure 3E) and acid-induced tolerance following adaptation (Figure 3F). Δ*glyA*, Δ*deoC*, Δ*acpP*, and Δ*rsgA* exhibited significantly reduced survival after 10 min of lethal acid shock in two assays (*P* < 0.05, Figure 3E and 3F), which indicated the mutants were more sensitive to acidic conditions. Of note, Δ*glyA* and Δ*acpP* showed lower viability after the pH 5.0 adaptation, whereas Δ*deoC* and Δ*rsgA* showed higher viability, suggesting that these genes may influence distinct facets of acid tolerance. Lastly, confocal imaging of 48 h biofilms revealed a higher fraction of dead cells in all four mutants compared with the wild type (Figure 3G and 3H), consistent with impaired fitness under acidic biofilm conditions. Collectively, these results demonstrate that the screen identified genes contributing to acid fitness across diverse physiological contexts, including growth, survival, and biofilm formation.

### 3.3 Genome-scale identification of oxidative stress tolerance determinants

Excessive oxidative stress can severely impair the viability of *S. mutans*. To identify genes required for oxidative stress tolerance, we applied 0.003% (*v*/*v*) H□O□, which represented the highest concentration that permitted measurable growth under our experimental conditions. CRISPRi sequencing was then performed to compare sgRNA abundance under normal (0% H_2_O_2_, CDY medium +1% xylose) and oxidative stress conditions (0.003% H_2_O_2_, CDY medium +1% xylose) (Figure 4A). Notably, a total of 337 significantly depleted genes were identified following H□O□ intervention (Table 2).

**Table 2.**
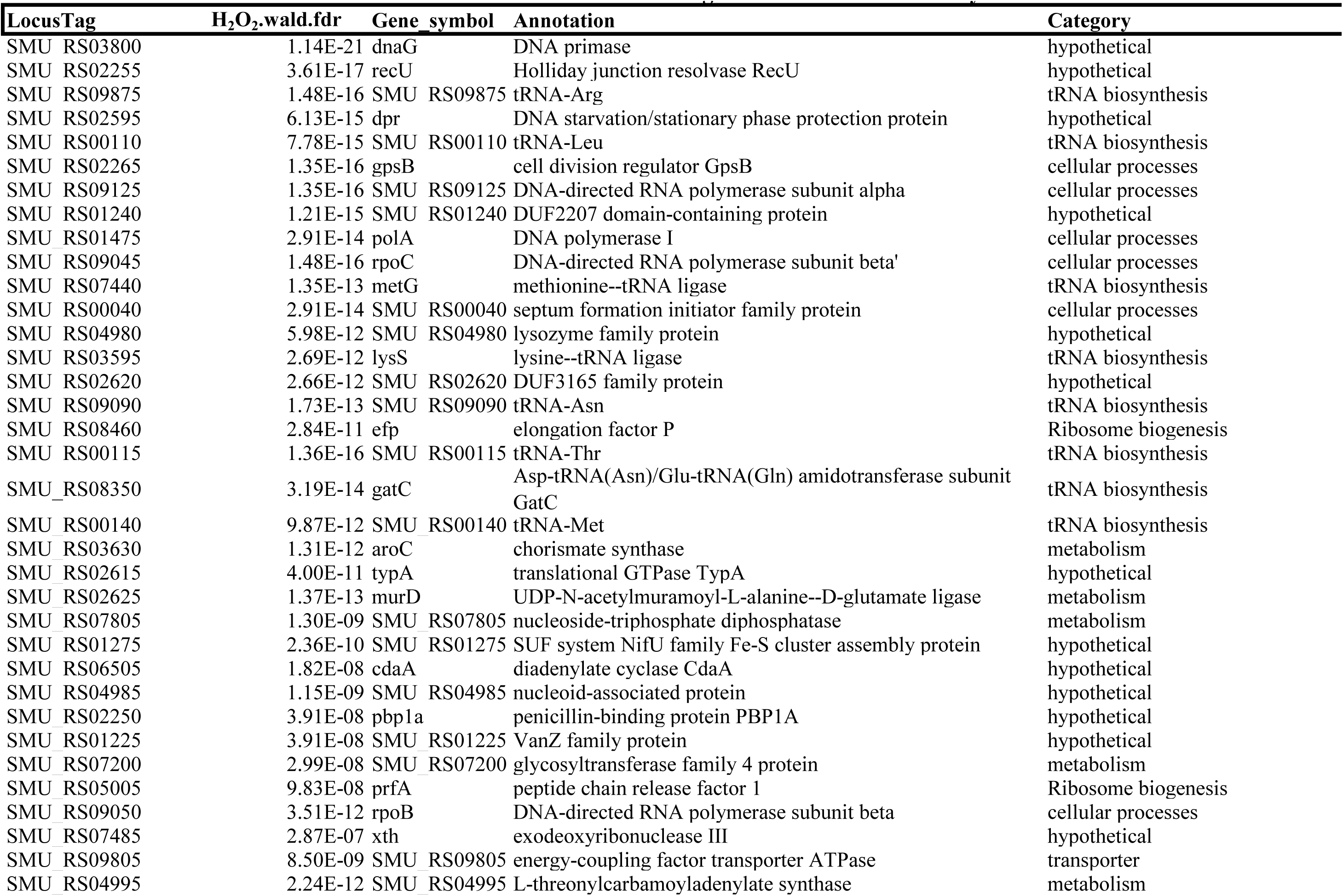
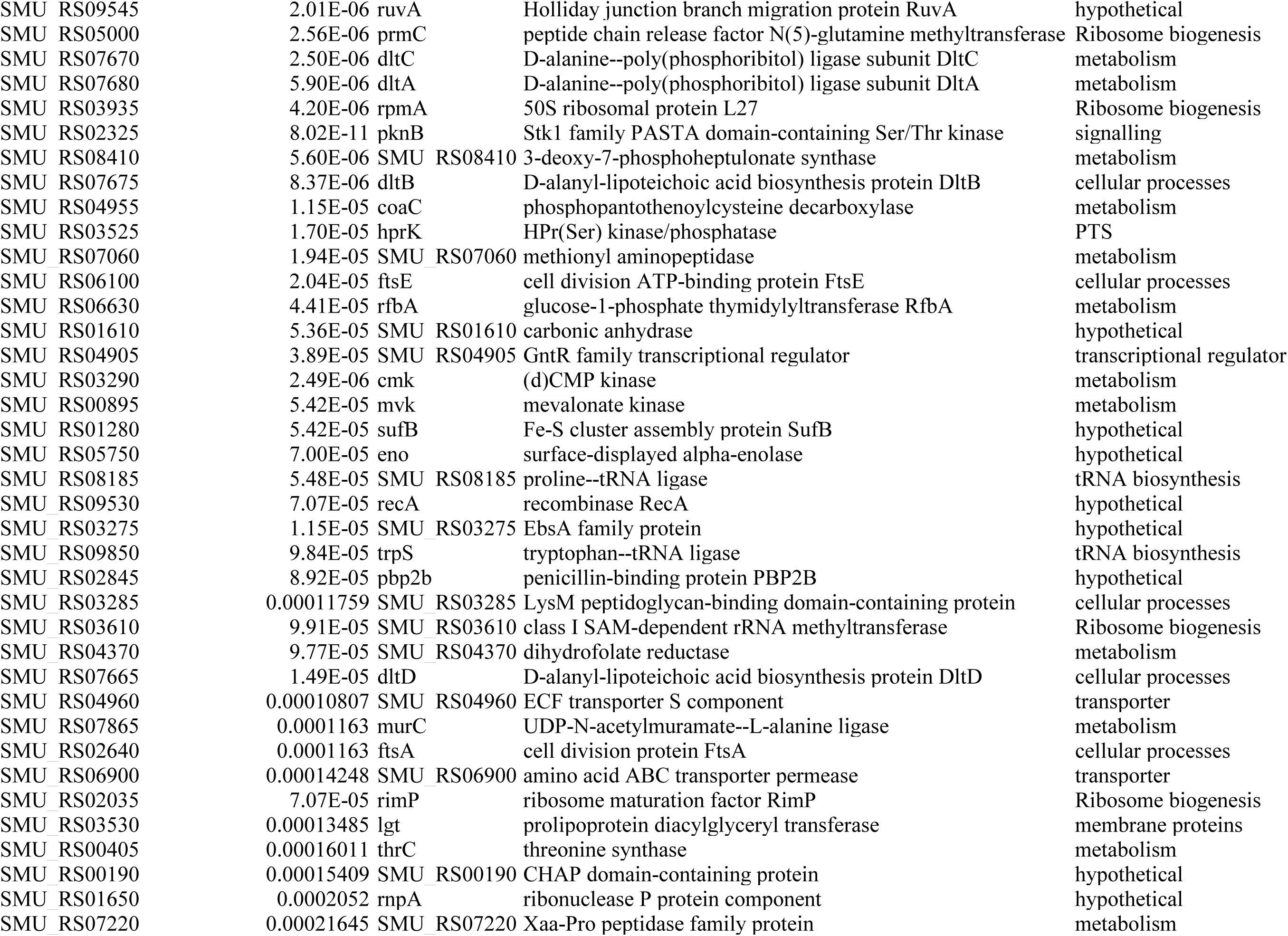
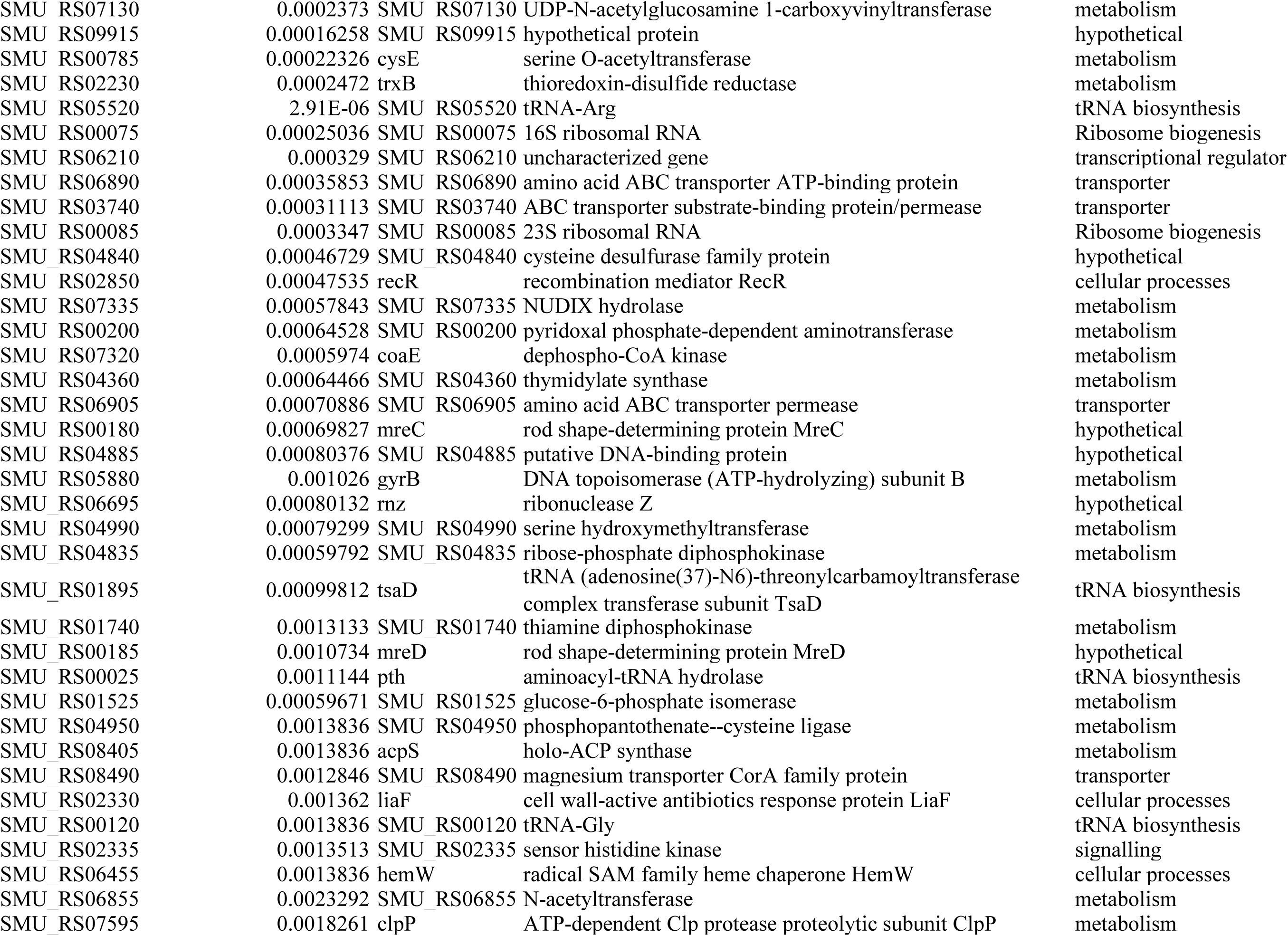
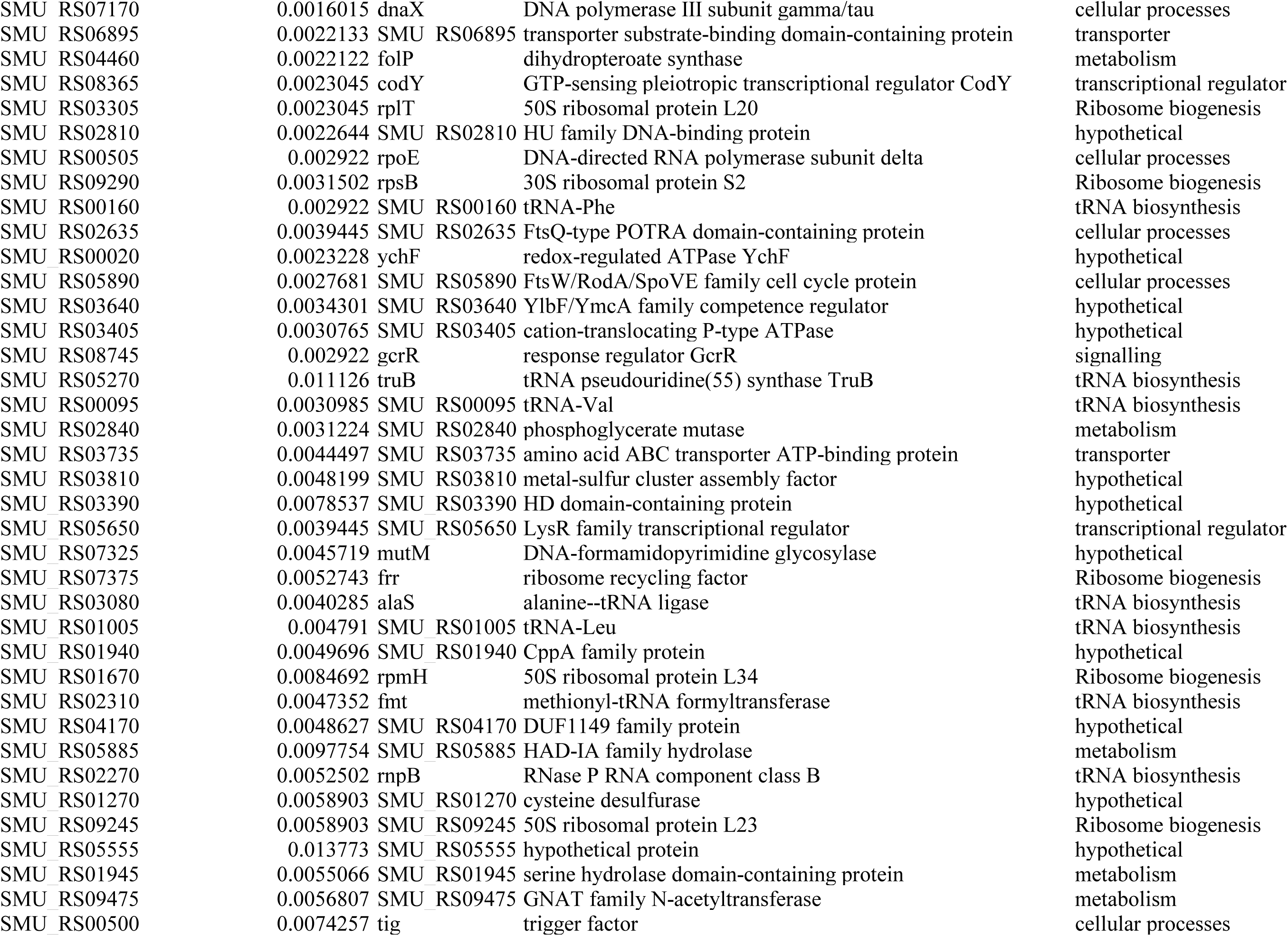
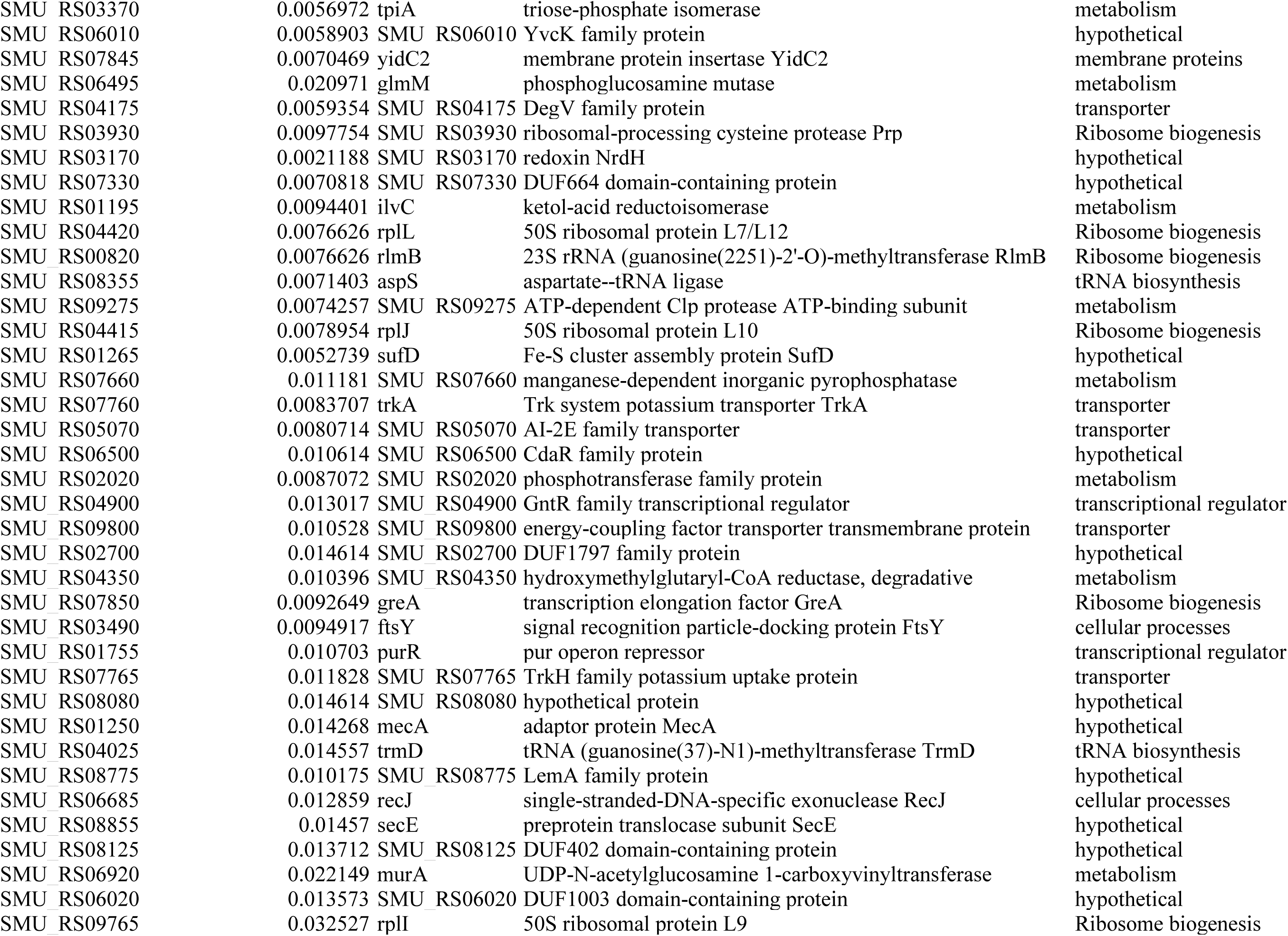
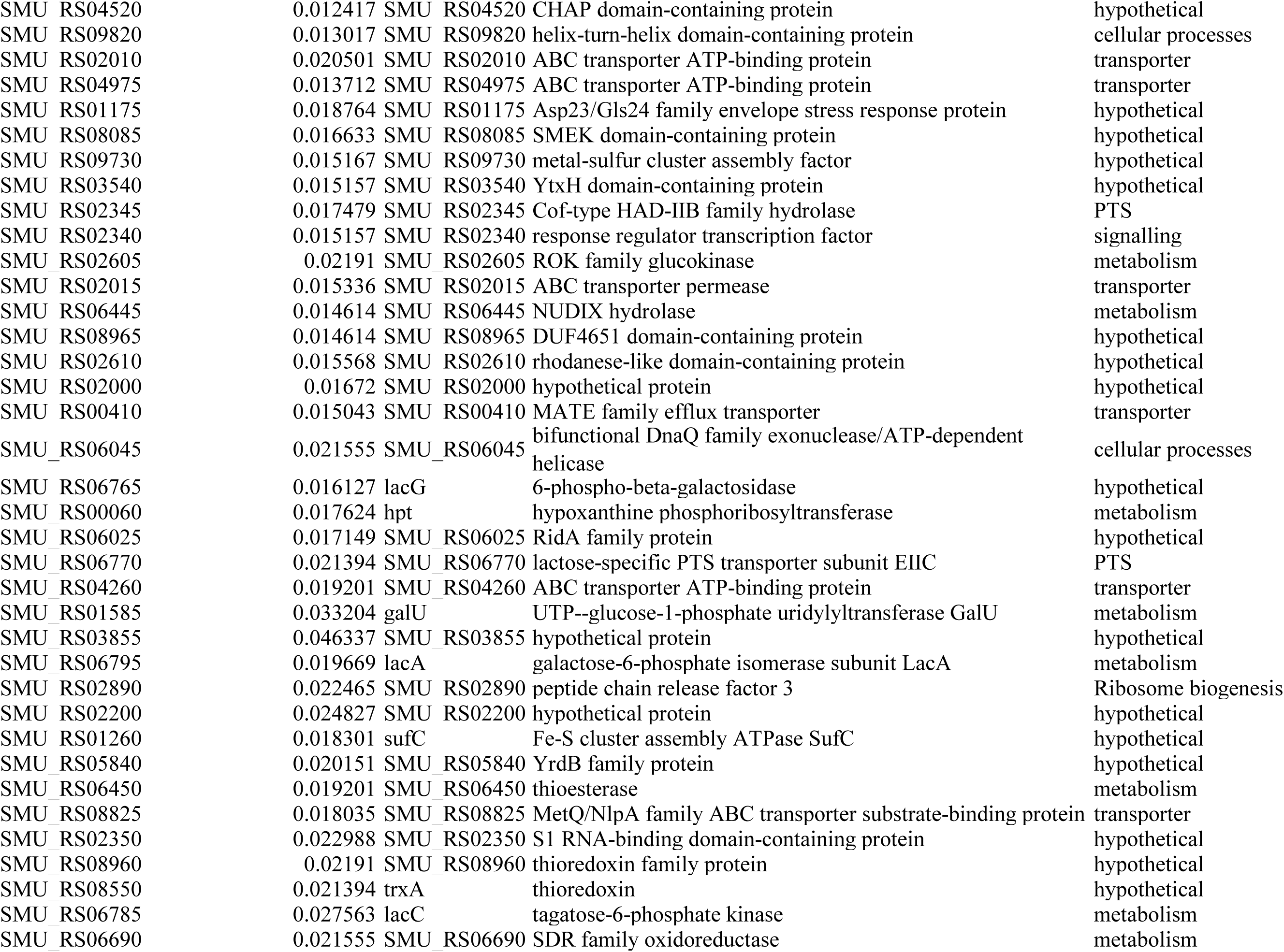
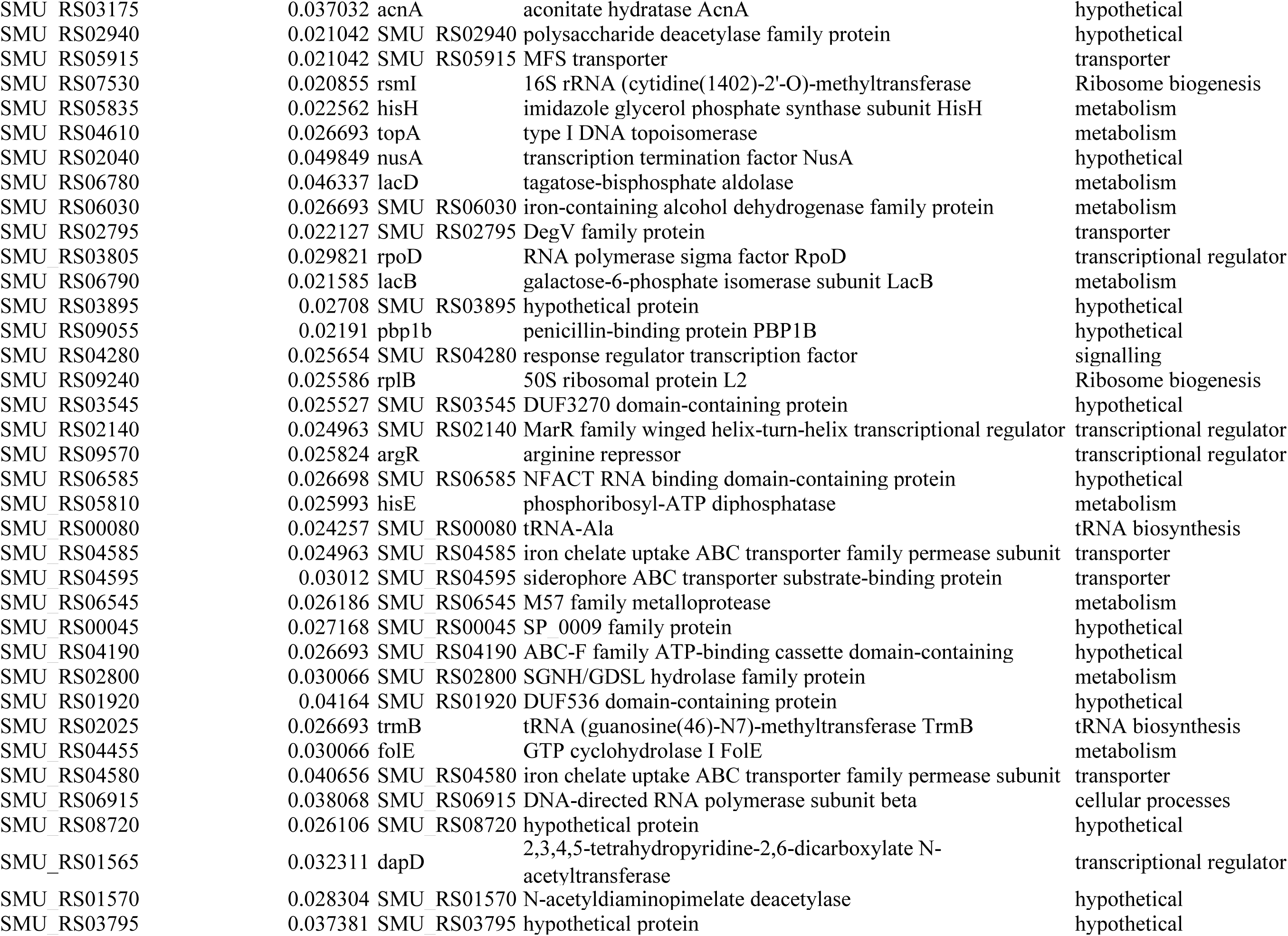
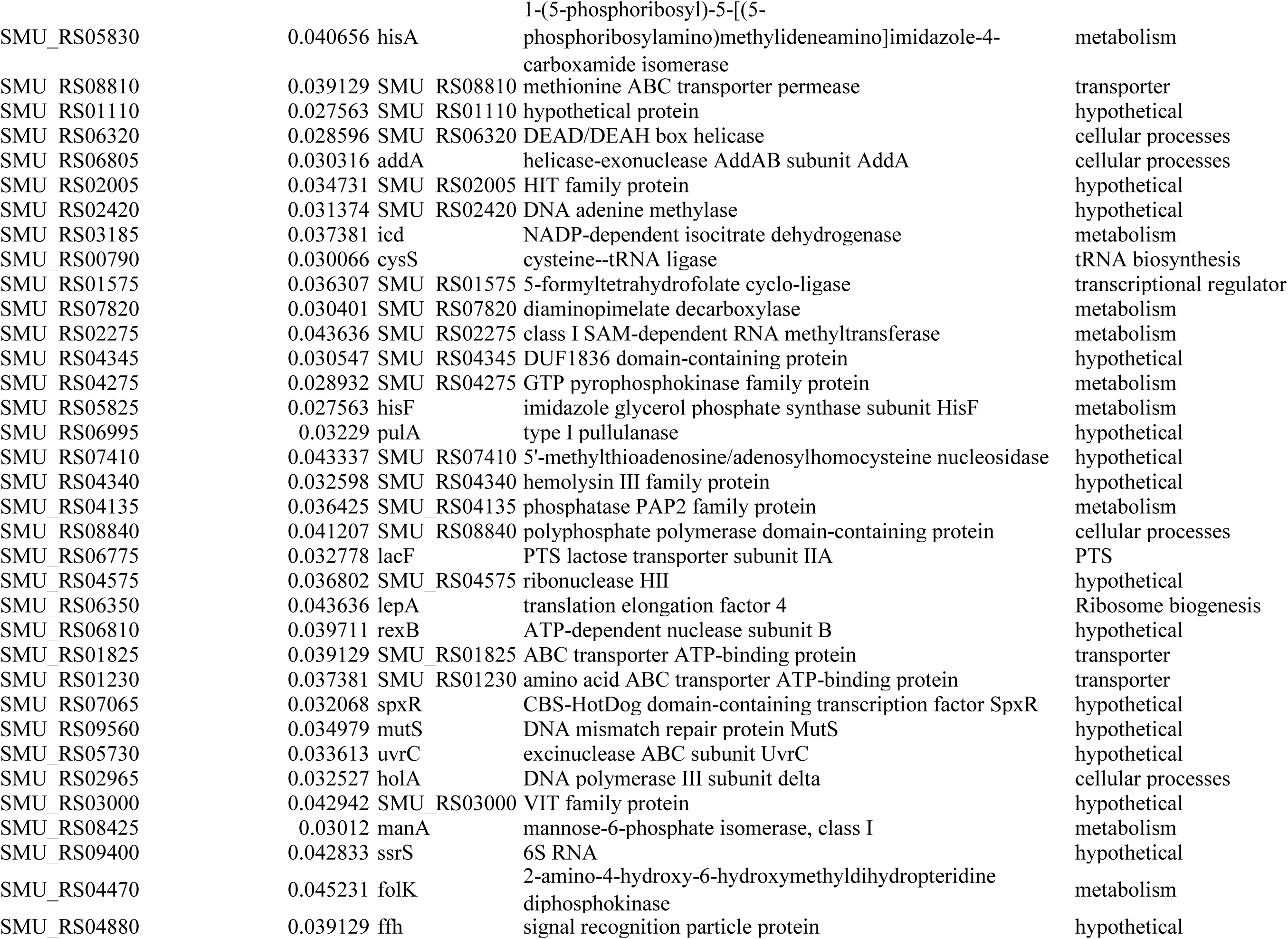
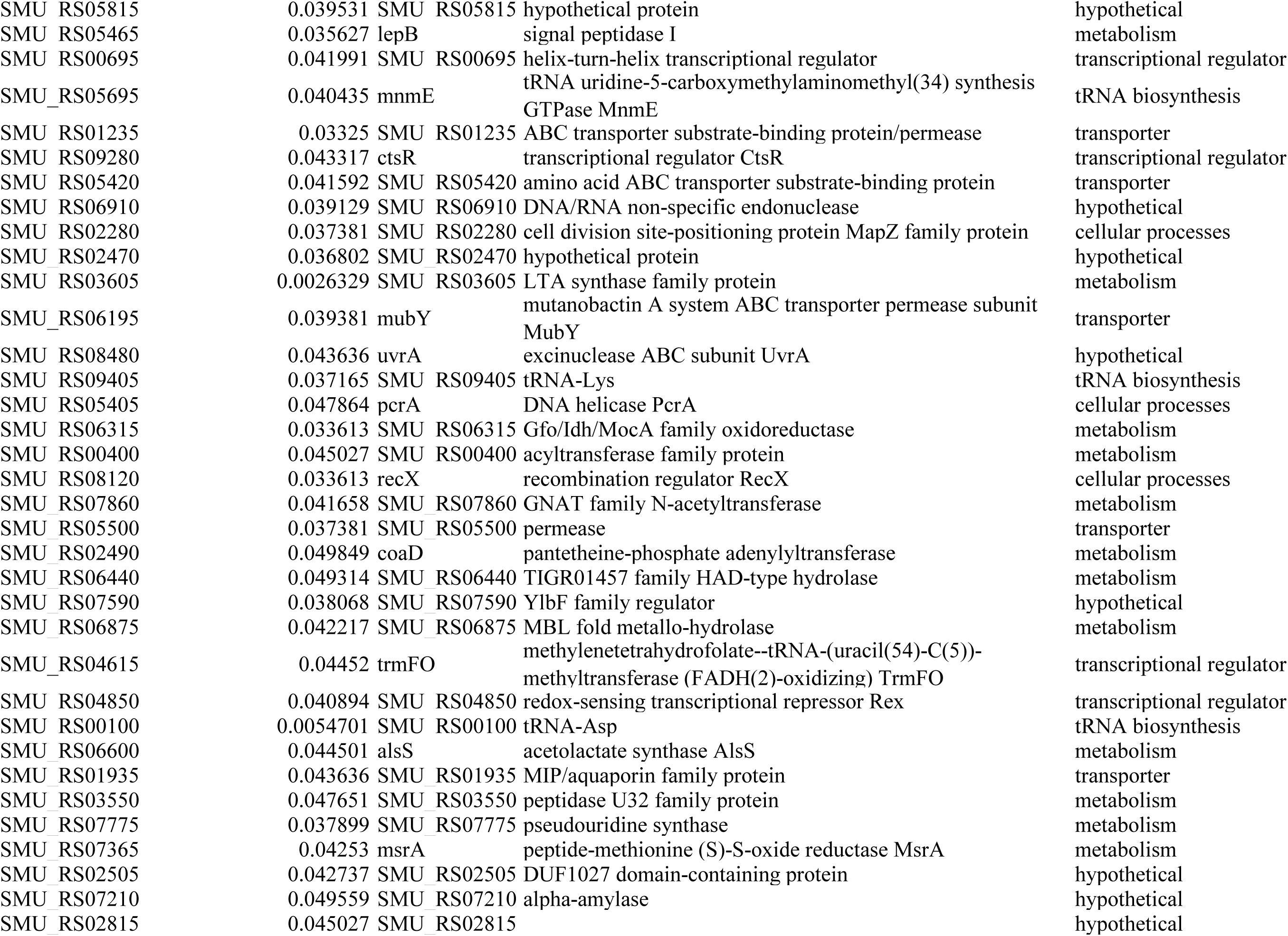
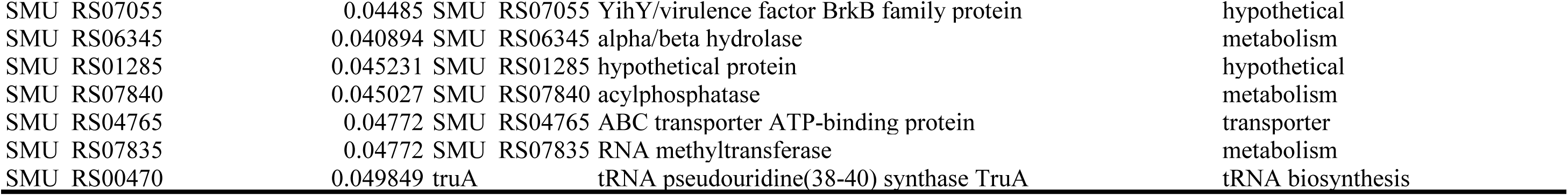
Oxidative stress tolerance genes identified in this study.

Moreover, GO enrichment analysis of these genes revealed functional features with both commonalities and differences compared to the acid tolerance response, suggesting key regulatory characteristics of *S. mutans* under oxidative stress (Figure 4C). At the BP level, macromolecule metabolic process (17.0%), nucleic acid metabolic process (14.0%), and macromolecule biosynthetic process (8.1%) were enriched, along with response to stress (6.3%) and regulation of cell shape (4.8%). At the CC level, intracellular anatomical structures were prominent (35.8%), consistent with intracellular sites of damage and repair. At the MF level, catalytic activity (25.1%) and exonuclease activity (2.2%) were enriched, supporting contributions from repair and redox-associated enzymatic functions.

To further validate the screen, deletion mutants were constructed for three representative genes (Δ*glyA*, Δ*gcrR*, and Δ*efp*) and their oxidative stress phenotypes were assessed relative to the wild-type strain. Under 0.003% H□O□, all mutants exhibited impaired growth, characterized by delayed entry into exponential phase and reduced stationary-phase cell yields (Figure 4D). In the inhibition zone assay, most gene-deficient strains showed no obvious difference in sensitivity compared to the wild-type under 0.1% (29 mM) H□O□ treatment, with only Δ*efp* displaying increased susceptibility. When the concentration was raised to 0.2% (58 mM), all tested deletion mutants displayed significantly larger inhibition zones than the wild-type, indicating their enhanced sensitivity to high-level oxidative stress (Figure 4E and 4F). Then, a hydrogen peroxide killing assay was conducted to quantify survival under oxidative challenge. After exposure to 0.01% H□O□ for 30 min, all gene-deficient strains showed markedly reduced viability relative to the wild-type (Figure 4G and 4H), consistent with observations from the inhibition zone assay. Together, these results demonstrate that the identified genes are required for resistance to oxidative stress in *Streptococcus mutans*.

### 3.4 Network organization and shared versus stress-specific genetic programs

To explore the functional connectivity among hits, a PPI network was constructed using the STRING database. For acid tolerance genes, 404 of the 422 depleted genes were connected in the inferred network (Figure 5A). This PPI network exhibited scale-free properties, with a clustering coefficient of 0.419 and significant enrichment relative to random expectation (P = 0.00295), suggesting biological organization rather than incidental connectivity. Hub proteins (Table S4) were further ranked in Cytoscape, with the highest-scoring hubs enriched in ribosome assembly and translation (rplI, rplJ, rplL, ffh, ftsY, rs6, rsfS), nucleic acid synthesis (purA, purC, pyrB, pyrF, pyrE), membrane protein assembly (yidC1, yidC2), stress-associated repair and proteostasis (dnaK, rpmA, ropA). Next, MCODE was applied to identify densely connected modules, detecting 26 functional clusters (Table S5). These modules were highly enriched for pathways involved in energy metabolism and carbon utilization, membrane transport and signal transduction, nucleic acid synthesis and repair, as well as ribosome assembly and translation regulation.

**Figure 5.**
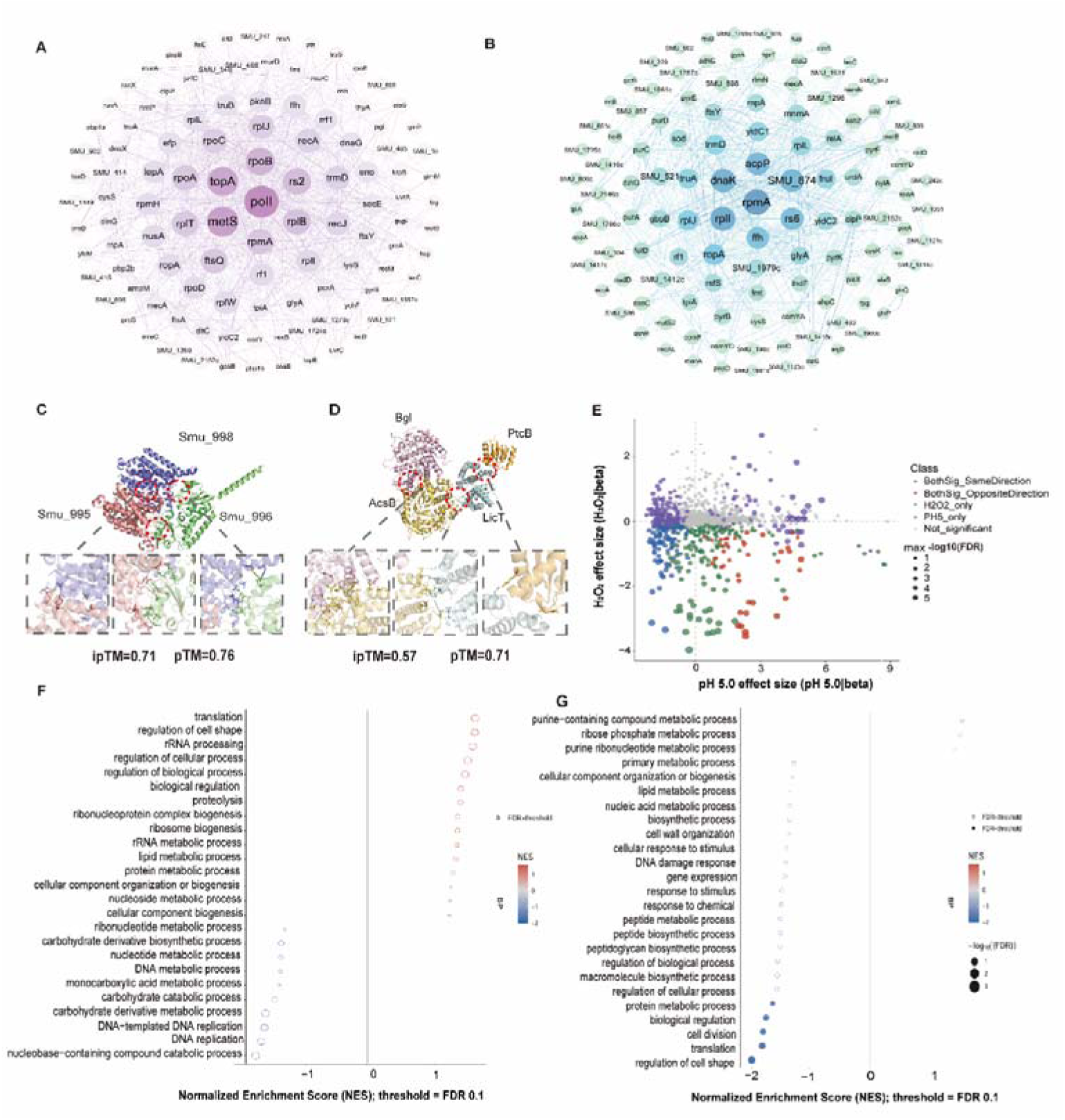
Integrated network, interaction, and functional analysis of acid and oxidative stress tolerance genes. (A, B) PPI network of acid-tolerant genes and oxidative stress tolerance genes generated by the STRING database and analyzed using Cytoscape, with hub genes defined as those having a degree value ≥ 10. (C) Predicted structures of oxidative stress-related gene modules (*smu_995*, *smu_996*, *smu_998*) and acid stress-related gene modules (*bgl*, *ptcB*, *acsB*, *licT*), where hydrogen bonds and salt bridges are formed on the lateral and longitudinal interfaces, with the distance between two amino acids analyzed by PDBePISA (http://www.ebi.ac.uk/pdbe/pisa). (E) Scatter plot comparing gene effect sizes (beta values) from CRISPRi screening under H□O□ (y-axis) and pH 5.0 (x-axis) stress. Genes are colored by their significance and directional consistency across conditions: blue (significant in both, same direction), red (significant in both, opposite direction), green (significant only in H□O□), purple (significant only in pH 5.0), and gray (non-significant). Point size indicates the maximum -log□□(FDR) value for each gene, reflecting statistical confidence. (F, G) Pre-ranked gene set enrichment analysis (GSEA) results showing biological process terms for concordant trends (F) and opposite trends (G). Circle size represents -log□□(FDR), and color indicates the direction of enrichment (blue = negatively enriched, red = positively enriched).

For oxidative stress tolerance genes, 317 of the 337 depleted genes formed interaction relationships (Figure 5B). The inferred network also exhibited scale-free properties, with a clustering coefficient of 0.393 and strong enrichment (*P* = 1.52 × 10^−13^), indicating non-random organization. Hubs were again emphasized in ribosome assembly (*rplJ* and *rplI*), transcription and translation regulation (*rpoA* and *efp*), translation termination and factor activity (*rf1* and *prfC*), protein secretion and localization, and protein folding and stress repair. MCODE identified 18 functional modules (Table S7); Module 1 comprised the largest core cluster and contained many key factors involved in translation and transcription, whereas additional modules were enriched for cell envelope biogenesis, energy metabolism, signal transduction, and metal ion homeostasis.

To assess module plausibility, we performed AlphaFold-based protein structure inspection of representative protein pairs within selected modules (Figure 5C and 5D). Although some modules displayed high-confidence structural interaction interfaces supporting physical association, others showed relatively lower structural confidence (Figure S3). Overall, these analyses provided partial structural support for the inferred modular organization, lending additional confidence to the functional clustering of genes associated with acid tolerance and H□O□ tolerance.

Finally, to delineate shared and stress-specific genetic programs, a combined analysis of the two differential-depletion datasets was conducted (Figure 5E). A total of 109 genes with concordant trends across both stresses and 43 genes with opposite trends were identified. Pre-ranked GSEA of the concordant gene set identified trends in translation, ribosome biogenesis, rRNA processing, DNA replication, and basal metabolism, although no biological process term reached FDR ≤ 0.1 and the overall signal was modest and dispersed (Figure 5F). Conversely, GSEA of the oppositely regulated genes identified five significantly enriched pathways (FDR < 0.1): protein metabolic process, biological regulation, cell division, translation, and regulation of cell shape (Figure 5G). These pathways were downregulated under H□O□ stress but upregulated under acid stress, consistent with distinct stress-dependent deployment of core growth-associated processes. Taken together, these findings suggest directional divergence and a resource-allocation trade-off in core growth-related processes between oxidative and acid stress responses.

## Discussion

Acid and oxidative stress tolerance are central to the cariogenic fitness of *S. mutans*. Yet most mechanistic models have focused on individual pathways or regulators rather than the broader genetic architecture underlying stress adaptation^[10]^. Herein, by establishing a genome-wide pooled CRISPRi platform, we provide a comparative functional map of genes that support fitness under acidic and HlJOlJ stress. Our findings highlight that the environmental stress adaptation of *S. mutans* does not depend on discrete, stress-specific pathways alone, but fundamentally relies on the integrity and stability of core physiological functional modules. Furthermore, we surmise that *S. mutans* adopts a precise resource allocation trade-off strategy to differentially tune core physiological pathways, enabling adaptive fitness under distinct stress conditions (Figure 6).

**Figure 6.**
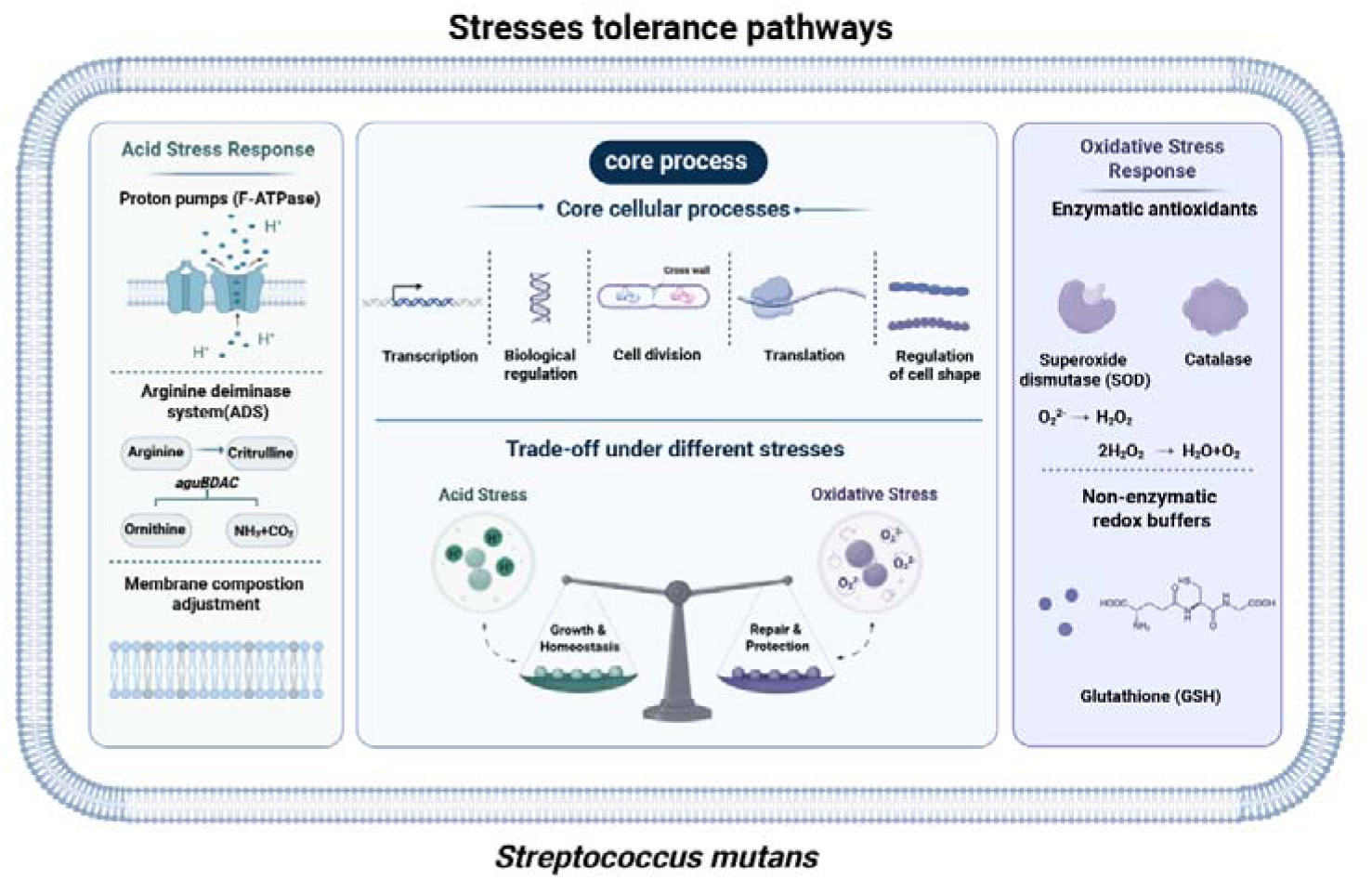
Summary of stress tolerance pathways in *S. mutans*. Rather than relying solely on dedicated stress-specific defense pathways, the adaptation of *S. mutans* to environmental stress fundamentally depends on the integrity and stability of conserved core physiological processes, including transcription, biological regulation, cell division, translation, and cell-shape maintenance. Under distinct environmental challenges, *S. mutans* reallocates cellular resources by differentially tuning these core physiological modules while activating stress-specific protective mechanisms. During acidic stress, cellular investment is preferentially directed toward growth- and homeostasis-associated functions, including proton extrusion, alkali generation, and membrane adaptation. In contrast, oxidative stress promotes resource allocation toward repair and protection through antioxidant enzymes and redox-buffering systems. This coordinated trade-off strategy enables *S. mutans* to optimize physiological fitness and maintain survival across diverse environmental conditions.

After excluding genes that significantly impaired the growth of *S. mutans*, we identified 422 genes specifically associated with acid tolerance. Many of the strongest signals mapped to fundamental processes that maintain cellular homeostasis, including DNA repair, RNA and tRNA metabolism, protein quality control, and envelope-associated functions. Notably, several genes belonging to canonical acid-tolerance pathways did not meet our final hit criteria. One plausible explanation is that some classical pathways impose large fitness costs even under non-stress conditions when partially repressed. Under our analytical framework, such genes are classified as general growth determinants rather than acid-specific fitness factors. For instance, the proton-translocating F_1_F_0_-ATPase is a canonical acid-resistance system, and several ATP synthase subunits showed strong acid-associated depletion, with large drop-fold values^[12, 37^]. Nevertheless, these genes were excluded from the dataset because their repression also caused major fitness defects under normal growth conditions. Consequently, our dataset is enriched for functions that become limiting specifically during acid challenge, rather than for universally required growth processes.

Collectively, these findings support a broader view of acid adaptation in *S. mutans*. Acid tolerance appears to be governed not only by dedicated acid-resistance circuits, but also by the robustness of core physiological systems that sustain growth while limiting and repairing damage. Within this framework, disruption of fundamental processes, including translation-associated functions, RNA/tRNA handling, protein quality control, and envelope homeostasis, can render cells highly susceptible to acid stress, even when canonical acid-resistance pathways remain intact. This interpretation aligns with our previous work on the cytoskeletal protein FtsZ, where we found that the *S. mutans* ortholog maintains function more effectively under acidic conditions than counterparts from other bacteria^[19, 38^].

Importantly, a similar pattern emerged in our oxidative stress screening. *S. mutans* is commonly thought to counter oxidative damage through detoxification and redox-control systems. However, under the low-dose H□O□ condition used here, only a subset of redox-related genes, such as *dpr, trxA, trxB,* and *rex*^[39, 40^], emerged as strong contributors to fitness. One possible explanation is that bacterial responses to H_2_O_2_ differ between resistance and tolerance states under distinct exposure regimes^[41]^. This distinction may partly account for the discrepancy between our findings and those of classical single-gene studies. It may also reflect the functional redundancy and buffering capacity of redox networks^[42]^, in which partial repression of one node does not necessarily produce a large pooled fitness defect^[20, 43^]. In this context, the prominence of *dpr, trxA, trxB,* and *rex* linked to iron homeostasis and thiol repair is consistent with a preventive strategy that limits secondary damage during prolonged oxidative exposure^[40]^. Beyond individual genes, network analysis provides a systems-level perspective on the organization of stress responses. Across both screens, identified hits were not randomly distributed but instead clustered into densely connected modules enriched for protein synthesis and maturation, nucleic acid metabolism, cell division, cell wall biogenesis, and membrane protein targeting. This modular organization supports a model in which stress tolerance is constrained by the capacity of core cellular systems to sustain growth while repairing stress-induced damage^[44]^.

In the present study, a comparative analysis of our dual-stress screening data uncovered a core adaptive strategy of *S. mutans* that tailors resource allocation among core physiological pathways to cope with distinct environmental stressors. Network module analysis demonstrated that hit genes from both acid and oxidative stress assays converged on overlapping core processes, including protein synthesis, nucleic acid metabolism, cell division, and cell wall biogenesis. Notably, some of these core pathways exhibited opposite regulatory patterns under the two stress conditions, representing canonical resource allocation trade-offs. In the chronically acidic and nutritionally fluctuating biofilm niche, *S. mutans* balances resource investment toward growth and stress survival. Sustained translation, metabolism, and cell division support robust proliferation, facilitating adaptation to chronic acid stress ^[45]^. In contrast, to survive, *S. mutans* reallocates metabolic resources by repressing growth programs and prioritizing damage repair, protein homeostasis, and membrane protection, thereby adopting a repair-first adaptive strategy^[46]^.

Nonetheless, our study has several limitations. First, hit calling depends on screening conditions and analysis thresholds, and these choices may bias discovery toward certain response modes^[47]^. Second, CRISPRi produces partial knockdown rather than complete loss of function, which can reduce sensitivity for buffered or redundant pathways^[43]^. Third, pooled screens report population-level fitness effects and may miss factors that act in spatially structured, multispecies biofilms^[48]^. Finally, network inferences and structural inspections provide supportive evidence, but they do not establish direct physical interactions or causal epistasis^[49, 50^]. Therefore, future work should test prioritized modules and hub genes using targeted genetic perturbations and epistasis designs, and evaluate key determinants in biofilm models that better approximate oral ecological complexity. Together, these efforts will refine the causal architecture of dual-stress adaptation and may inform strategies to disrupt cariogenic fitness.

## Supporting information

Supplymentry material PCR process of sgRNA sequencing

Supplymentry material pYL02 sequence

## CRediT authorship contribution statement

Yaqi Chi: Methodology, Investigation, Formal analysis, Writing – original draft. Yuxing Chen: Methodology, Investigation, Formal analysis, Writing – original draft. Chongyang Yuan: Methodology, Investigation, Formal analysis. Liuchang Yang: Investigation. Mingrui Zhang: Investigation. Xiaolin Chen: Investigation. Yiran Zhao: Investigation. Ming Li: Writing-review and editing, Supervision. Xiaoyan Wang: Resources, Funding acquisition, Project administration. Yongliang Li: Methodology, Investigation, Validation, Formal analysis, Conceptualization, Writing – review and editing, Supervision.

## Acknowledgements

This work was supported by the National Natural Science Foundation of China (Grant number: 82001039); the Fundamental Research Funds for the Central Universities and Young Elite Scientist Sponsorship Program by CAST (Grant number: 2019QNRC001 to YL.L). The Fundamental Research Funds for the Central Universities (PKU2024XGK001); The Open Research Fund Project of Key Laboratory of Stomatology of Hebei Province (Grant NO. HBKLS-KF202401) to X.Y. Wang; We also thank Dr. Jiao Liu from the Center of Medical and Health Analysis, Peking University Health Science Center, for assistance with confocal microscopy imaging.

