## Supplementary material for "Defining the genetic landscape of acid and oxidative stress tolerance in *Streptococcus mutans* by pooled CRISPR interference screening": Supplymentry material PCR process of sgRNA sequencing

**The primers and** **PCR procedure of sgRNA sequencing.**

1.1 **First round of PCR amplification**

**PCR amplification primers:**

| NGS-F | ACACTCTTTCCCTACACGACGCTCTTCCGATCTACATTGCACTGTCCCCCTGG |
| --- | --- |
| NGS-R | GACTGGAGTTCAGACGTGTGCTCTTCCGATCTATGCTGTTTCCAGCATAGCTC |

**PCR reaction components:**

| Reagent | Volume (μl) |
| --- | --- |
| ddH_2_O  5×PCR Buffer  10 mM dNTP  Forward primer (20 pmol/μl)  Reverse primer (20 pmol/μl)  Hot start polymerase  Template DNA | Up to 50  10  1  1  1  1  500ng |
| Total | 50 |

**PCR conditions:**

| Predenaturation 98℃ | 2 min |
| --- | --- |
| Denaturation 98℃  Annealing 58℃  Extension 72℃ | 30 s  30 s (15 cycles)  30s |
| Final extension 72℃  4℃ | 2min  ∞ |

**1.2 Second round of PCR amplification (using the PCR product from the first round of PCR as a template)**

**PCR amplification primers:**

| NGS-P5 | AATGATACGGCGACCACCGAGATCTACACTATAGCCTACACTCTTTCCCTACACGACGCTCTTCCGATCT |
| --- | --- |
| NGS-P7 | CAAGCAGAAGACGGCATACGAGATCGAGTAATGTGACTGGAGTTCAGACGTGTGCTCTTCCGATCT |

**PCR reaction components:**

| Reagent | Volume (μl) |
| --- | --- |
| ddH_2_O  5×PCR Buffer  10 mM dNTP  Forward primer (20 pmol/μl)  Reverse primer (20 pmol/μl)  Hot start polymerase  Product from the 1^st^ round of PCR | 34  10  1  1  1  1  2 |
| Total | 50 |

**PCR conditions:**

| Predenaturation 98℃ | 2 min |
| --- | --- |
| Denaturation 98℃  Annealing 58℃  Extension 72℃ | 30 s  30 s (20 cycles)  30s |
| Final extension 72℃  4℃ | 2min  ∞ |
