## Supplementary material for "Defining the genetic landscape of acid and oxidative stress tolerance in *Streptococcus mutans* by pooled CRISPR interference screening": Supplymentry material pYL02 sequence

acatgtgagcaaaaggccagcaaaaggccaggaaccgtaaaaaggccgcgttgctggcgtttttccataggctccgcccccctgacgagcatcacaaaaatcgacgctcaagtcagaggtggcgaaacccgacaggactataaagataccaggcgtttccccctggaagctccctcgtgcgctctcctgttccgaccctgccgcttaccggatacctgtccgcctttctcccttcgggaagcgtggcgctttctcatagctcacgctgtaggtatctcagttcggtgtaggtcgttcgctccaagctgggctgtgtgcacgaaccccccgttcagcccgaccgctgcgccttatccggtaactatcgtcttgagtccaacccggtaagacacgacttatcgccactggcagcagccactggtaacaggattagcagagcgaggtatgtaggcggtgctacagagttcttgaagtggtggcctaactacggctacactagaaggacagtatttggtatctgcgctctgctgaagccagttaccttcggaaaaagagttggtagctcttgatccggcaaacaaaccaccgctggtagcggtggtttttttgtttgcaagcagcagattacgcgcagaaaaaaaggatctcaagaagatcctttgatcttttctacggggtctgacgctcagtggaacgaaaactcacgttaagggattttggtcatgagattatcaaaaaggatcttcacctagatccttttaaattaaaaatgaagttttaaatcaatctaaagtatatatgagtaaacttggtctgacagttaccaatgcttaatcagtgaggcacctatctcagcgatctgtctatttcgttcatccatagttgcctgactccccgtcgtgtagataactacgatacgggagggcttaccatctggccccagtgctgcaatgataccgcggcttccacgctcaccggctccagatttatcagcaataaaccagccagccggaagggccgagcgcagaagtggtcctgcaactttatccgcctccatccagtctattaattgttgccgggaagctagagtaagtagttcgccagttaatagtttgcgcaacgttgttgccattgctacaggcatcgtggtgtcacgctcgtcgtttggtatggcttcattcagctccggttcccaacgatcaaggcgagttacatgatcccccatgttgtgcaaaaaagcggttagctccttcggtcctccgatcgttgtcagaagtaagttggccgcagtgttatcactcatggttatggcagcactgcataattctcttactgtcatgccatccgtaagatgcttttctgtgactggtgagtactcaaccaagtcattctgagaatagtgtatgcggcgaccgagttgctcttgcccggcgtcaatacgggataataccgcgccacatagcagaactttaaaagtgctcatcattggaaaacgttcttcggggcgaaaactctcaaggatcttaccgctgttgagatccagttcgatgtaacccactcgtgcacccaactgatcttcagcatcttttactttcaccagcgtttctgggtgagcaaaaacaggaaggcaaaatgccgcaaaaaagggaataagggcgacacggaaatgttgaatactcatactcttcctttttcaatattattgaagcatttatcagggttattgtctcatgagcggatacatatttgaatgtatttagaaaaataaacaaataggggttccgcgcacatttccccgaaaagtgccacctgacgtctaagaaaccattattatcatgacattaacctataaaaataggcgtatcacgaggccctttcgtctcgcgcgtttcggtgatgacggtgaaaacctctgacacatgcagctcccggagacggtcacagcttgtctgtaagcggatgccgggagcagacaagcccgtcagggcgcgtcagcgggtgttggcgggtgtcggggctggcttaactatgcggcatcagagcagattgtactgagagtgcaccataaaattgtaaacgttaatattttgttaaaattcgcgttaaatttttgttaaatcagctcattttttaaccaataggccgaaatcggcaaaatcccttataaatcaaaagaatagcccgagatagggttgagtgttgttccagtttggaacaagagtccactattaaagaacgtggactccaacgtcaaagggcgaaaaaccgtctatcagggcgatggcccactacgtgaaccatcacccaaatcaagttttttggggtcgaggtgccgtaaagcactaaatcggaaccctaaagggagcccccgatttagagcttgacggggaaagccggcgaacgtggcgagaaaggaagggaagaaagcgaaaggagcgggcgctagggcgctggcaagtgtagcggtcacgctgcgcgtaaccaccacacccgccgcgcttaatgcgccgctacagggcgcgtactatggttgctttgacgtatgcggtgtgaaataccgcacagatgcgtaaggagaaaataccgcatcaggcgcccctgcaggcagctgcgcgctcgctcgctcactgaggccgcccgggcaaagcccgggcgtcgggcgacctttggtcgcccggcctcagtgagcgagcgagcgcgcagagagggagtggccaactccatcactaggggttcctgcggccgcTCCCCAGCATGCCTGCTATTcTCTTCCCAATCCTCCCCCTTGCTGTCCTGCCCCACCCCACCCCCCAGAATAGAATGACACCTACTCAGACAATGCGATGCAATTTCCTCATTTTATTAGGAAAGGACAGTGGGAGTGGCACCTTCCAGGGTCAAGGAAGGCACGGGGGAGGGGCAAACAACAGATGGCTGGCAACTAGAAGGCACAGTCGAGGCTGATCAGCGAGCTCTAGgaattcttacttgtctagttcttccgctagtgaagtaaaagagattttatttgcacttattcatggttttgtaacccatatgcctgagtccccttatggtctagtcggaataggaatttgtgttccaggccttgtagatcgtcatcagcaaattattttcatgcctaacttaaattggaatatcaaagatttgcagtttttaattgagagtgagtttaatgttccggtttttgttgaaaatgaagctaatgcaggagcatacggtgaaaaagtatttggtatgacaaaaaactatgaaaacatcgtttacatcagtattaatatcggaattggaactggacttgttattaacaacgaattgtataaaggtgttcagggtttttctggggaaatgggtcatatgacgatagattttaatggacccaaatgcagctgtggaaatcgaggctgttgggaattatatgcttctgaaaaagcgttactggcttcgctctctaaagaagaaaagaatatttctcgaaaagagattgtggaacgcgcaaataaaaatgatgtagaaatgttaaatgcacttcaaaactttggcttttatatcggaattggattaaccaatatccttaatacatttgatatagaagctgttatcttgagaaatcatataattgaatctcatcccattgttttaaatacgattaaaaacgaagtttcttctagagtccattctcatttagacaataaatgtgaactattgccttcttcgttaggaaaaaatgCacctgctttaggagcggtttctatcgttattgattcttttttaagtgttacccctataagttagAGAAGATCGATTTTCGTTCGTGAATAAATAACGTAACGTGACTGGCAAGAGATATTTTTAAAACAATGAATAGGTTTACACTTACTTTAGTTTTATGGAAATGAAAGATCATATCATATATAATCTAGAATAAAATTAACTAAAATAATTATTATCTAGATAAAAAATTTAGAAGCCAATGAAATCTATAAATAAACTAAATTAAGTTTATTTAATTAACAACTATGGATATAAAATAGGTACTAATCAAAATAGTGAGGAGGATATATTTGAATACATACGAACAAATTAATAAAGTGAAAAAAATACTTCGGAAACATTTAAAAAATAACCTTATTGGTACTTACATGTTTGGATCAGGAGTTGAGAGTGGAaattttgtcaaaataattttattgacaacgtcttattaacgttgatataatttaaattttatttccattctacagtttattcttgacattgcactgtccccctggtataataacatgtggGTCTTCgaGAAGACctgtttaagagctatgctggaaacagcatagcaagtttaaataaggctagtccgttatcaacttgaaaaagtggcaccgagtcggtgcttttttctgtctcttatacacatctgacgctgccgacgaTTTTTatgacaaaaagagaaaattttgataaaatagtcttattaactaataaggaggacaaacATGAACAAAAATATAAAATATTCTCAAAACTTTTTAACGAGTGAAAAAGTACTCAACCAAATAATAAAACAATTGAATTTAAAAGAAACCGATACCGTTTACGAAATTGGAACAGGTAAAGGGCATTTAACGACGAAACTGGCTAAAATAAGTAAACAGGTAACGTCTATTGAATTAGACAGTCATCTATTCAACTTATCGTCAGAAAAATTAAAACTGAATACTCGTGTCACTTTAATTCACCAAGATATTCTACAGTTTCAATTCCCTAACAAACAGAGGTATAAAATTGTTGGGAGTATTCCTTACCATTTAAGCACACAAATTATTAAAAAAGTGGTTTTTGAAAGCCATGCGTCTGACATCTATCTGATTGTTGAAGAAGGATTCTACAAGCGTACCTTGGATATTCACCGAACACTAGGGTTGCTCTTGCACACTCAAGTCTCGATTCAGCAATTGCTTAAGCTGCCAGCGGAATGCTTTCATCCTAAACCAAAAGTAAACAGTGTCTTAATAAAACTTACCCGCCATACCACAGATGTTCCAGATAAATATTGGAAGCTATATACGTACTTTGTTTCAAAATGGGTCAATCGAGAATATCGTCAACTGTTTACTAAAAATCAGTTTCATCAAGCAATGAAACACGCCAAAGTAAACAATTTAAGTACCGTTACTTATGAGCAAGTATTGTCTATTTTTAATAGTTATCTATTATTTAACGGGAGGAAATAAccagtgttgaatctttagattaagggaaactcttataattaaaagaactagctcttgttctgcgttttctataagttagacagtatacttaactcatagagtcagtacaagcagctagctttttgttgtgataagacttgccaactgataaaaactcaggtaaactgttaatggtaatcaaaataaaaggagtttatatgtctttacctaattgtcctaaatgtcagtcagagtatgtctatgaagatggtattctattggtttgtccagaatgtgcttatgaatggaatcctgcagaagttgagaaagaagaaggacttgttgttattgatgcgaatggcaaacaattagctgatggtgatacagtcactcttattaaggatcttaaagttaagggggctcctaaagatttaaaacaaggaacacgtgttaaaaatattcgccttgttgaaggagatcacaatattgattgtaaaattgatggctttggtgctatgaaattaaagtctgaatttgtgaaaaaattataagagtagagcaagttgaactgtctatcttagaaacaatcaattaatgaaggtggttcattcttgttcttttttggtgatacaggaatctctgcttggttttatgcttatctttgaatgaaaattcaattccatctataaaattttctgaaaatttgatataataaaaacgattattttttgaggagcaagtatgtcttatatatcagaagttttaccgagtcttttagatggggcgctgattactttacaagtcttttttattgttattctcttttcaattcctttaggagctatattagctttccttatgcaggtaccttttagacctttacgttggctgttaaatctctatgtttggattatgcgtgggacaccgctccttttacaactgatttttatttattatgttttgccaagtgcagggatcacttttgatcgtatgccagcagctattttagcttttactctcaactatgctgcctatatgaaggaatcatgggaaataggccctc
