## Supplementary material for "Defining the genetic landscape of acid and oxidative stress tolerance in *Streptococcus mutans* by pooled CRISPR interference screening": Supplymentry material legend

**Figure S1 Rationale for 0.003% hydrogen peroxide screening condition**

(A) Bacterial culture images after 48‑h growth under 0.3%, 0.03%, 0.003%, 0.0003%, and 0.00003% H₂O₂ treatments, showing the minimum inhibitory concentration (MIC) lies between 0.003% and 0.03%. (B) Growth curves of bacteria cultured for 24 h under 0%, 0.0015%, 0.003%, and 0.006% H₂O₂.

**Figure S2 Characterization of the optimized genome-wide sgRNA library.**
(A) Relative repression activity of 1,919 sgRNAs selected from 49,694 candidates after off-target evaluation, plotted against their relative distance to the start codon. Nearly all selected sgRNAs showed approximately 100% predicted repression activity. (B) Frequency distribution of the relative distances between the selected sgRNAs and their corresponding start codons.

**Figure S3 AlphaFold3 structural prediction of selected protein-protein interaction modules in *Streptococcus mutans*.**

(A) A high-confidence interaction module (comprising Smu_1118c, Smu_1119c, Smu_1121c, Cdd, and DeoC) predicted by AlphaFold3. The complex shows a well-defined interaction interface supported by high ipTM (0.57) and pTM (0.65) scores, with detailed insets highlighting predicted hydrogen bonds and salt bridges at the protein-protein interfaces, indicative of stable physical association.

(B) A representative low-confidence interaction module (comprising Smu_414, RexB, and Smu_393). The complex exhibits a lower ipTM score (0.23), indicating reduced structural confidence in the predicted interaction interface. While the overall pTM score (0.6) suggests plausible folding of individual proteins, the interaction interface lacks well-defined contacts, reflecting uncertainty in the physical association between these proteins.
