## Supplementary material for "Defining the genetic landscape of acid and oxidative stress tolerance in *Streptococcus mutans* by pooled CRISPR interference screening": Table S1

Table S1 Bacterial strains and plasmid used in this study

| **Strain or plasmid** | **Description** | **Source of reference** |
| --- | --- | --- |
| UA159 | *S. mutans* UA159 | ATCC 700610 |
| *ΔglyA* | UA159; *ΔglyA* :: IFDC2, Erm^R^ | This study |
| *ΔdeoC* | UA159; *ΔdeoC* :: IFDC2, Erm^R^ | This study |
| *ΔacpP* | UA159; *ΔacpP* :: IFDC2, Erm^R^ | This study |
| *ΔrsgA* | UA159; *ΔrsgA* :: IFDC2, Erm^R^ | This study |
| *Δefp* | UA159; *Δefp* :: IFDC2, Erm^R^ | This study |
| *ΔgcrR* | UA159; *ΔgcrR* :: IFDC2, Erm^R^ | This study |
| *ftsW-CRISPRi* | UA159/pUC19-P3-sgRNA-*ftsW*, Ermᴿ | This study |
| *pbp2x-CRISPRi* | UA159/pUC19-P3-sgRNA-*pbp2x*, Ermᴿ | This study |
| pYL02 | UA159; pUC19::P3-sgRNA, Erm^R^ | This study |
