## Supplementary material for "Defining the genetic landscape of acid and oxidative stress tolerance in *Streptococcus mutans* by pooled CRISPR interference screening": Table S2

Table S2 Primers used in PCR-ligation mutagenesis

| **Primer** | **Nucleotide sequence** |
| --- | --- |
| IFDC2-F | CCGAGCAACAATAACACTCATAGCAT |
| IFDC2-R | CGTCCCTTTAGTAACGTGTAACTTTCCAA |
| up-*acpP*-F | ATGAAGAAAATTGCAGTTGATGCTATGGG |
| up-*acpP*-R | ATGCTATGAGTGTTATTGTTGCTCGGCTTAGTCCCCCCTGTTTGAAAATTCT |
| dn-*acpP*-F | TTGGAAAGTTACACGTTACTAAAGGGACGATAAATAATACTAATGTTGTTAATAATGGAGGGTTTAGT |
| dn-*acpP*-R | AATCACATTTTCGTCTTCTGTCGTCAAAA |
| up-*deoC*-F | AGCCAGTTTTGGAGTGCCG |
| up-*deoC*-R | TTGGAAAGTTACACGTTACTAAAGGGACGGTTATCCTTTCTTAATAATCAAGTAATGATTTTTAAAATTTCAG |
| dn-*deoC*-F | ATGCTATGAGTGTTATTGTTGCTCGGTTTAATCAGAACAGCTATTAAAGCTAGTAAAAATGCT |
| dn-*deoC*-R | TTCAAAACGAGTGATCACTTCAGATTTCACA |
| up-*glyA*-F | TCAGAAGATGCTCTTAATTTAGCACAGGA |
| up-*glyA*-R | GGAAAGTTACACGTTACTAAAGGGACGTTTTTCTCCTTAAAATCTTATATTTTTCACATGCGC |
| dn-*glyA*-F | ATGCTATGAGTGTTATTGTTGCTCGGTGGATTTGTATTTAAAAAAGATTGTCATTCACCAATTTACT |
| dn-*glyA*-R | TGATTCGTATTAGTTTTTTTTTCATATTATTTACTTTGTATGTCTTCG |
| up-*rsgA*-F | GCCAGATTGGCAGATGCTCT |
| up-*rsgA*-R | ATGCTATGAGTGTTATTGTTGCTCGGACGCTCTCCTTACTAAACTTAATAGTTTCATT |
| dn-*rsgA*-F | TTGGAAAGTTACACGTTACTAAAGGGACGGTCTGTTATGTTGCTCAATCAAATTGCC |
| dn-*rsgA*-R | TCGCTAGGTAAAAAAATATTGGACAGCATATG |
| up-*efp*-F | TCCTTACCTAATTAATGAGCATCAGTTCAATGT |
| up-*efp*-R | ATGCTATGAGTGTTATTGTTGCTCGGGGAAAACAGTTTTGGATGAATTTATTAGAAAGGACA |
| dn-*efp*-F | TGGAAAGTTACACGTTACTAAAGGGACGTTTTTACTATACCTCTTTATAAAATATTATCATC |
| dn-*efp*-R | CTATAATACAGCTTATCGTATTCATCCCTTTGCT |
| up-*gcrR*-F | CTTCTTTTCACCGTTATCAAAAAAACTGACAAAAT |
| up-*gcrR*-R | ATGCTATGAGTGTTATTGTTGCTCGGATGTCTAGGTCTGTTGAAGTGGTTCA |
| dn-*gcrR*-F | TGGAAAGTTACACGTTACTAAAGGGACGATACTCCTCAACAAAACTCTAACAATTTCT |
| dn-*gcrR*-R | CTATACTTGGTTTGATGAAGAAACTCCACGG |
