## Supplementary figures and images for "Defining the genetic landscape of acid and oxidative stress tolerance in *Streptococcus mutans* by pooled CRISPR interference screening"

### Figure S1

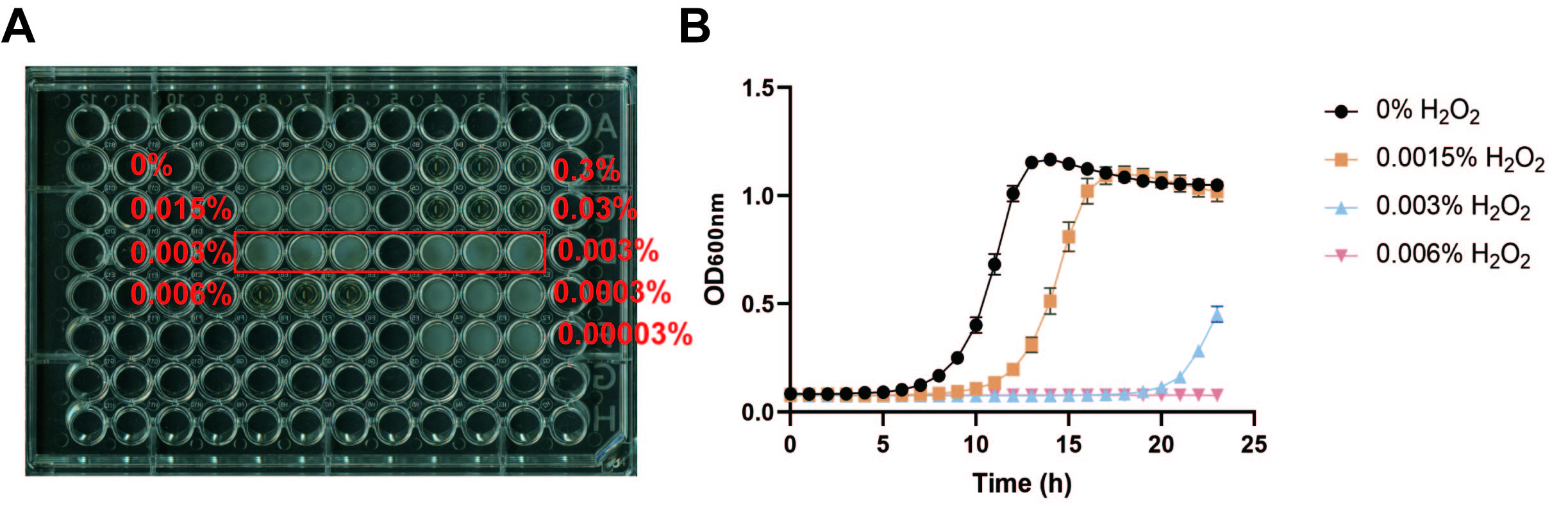

### Figure S2

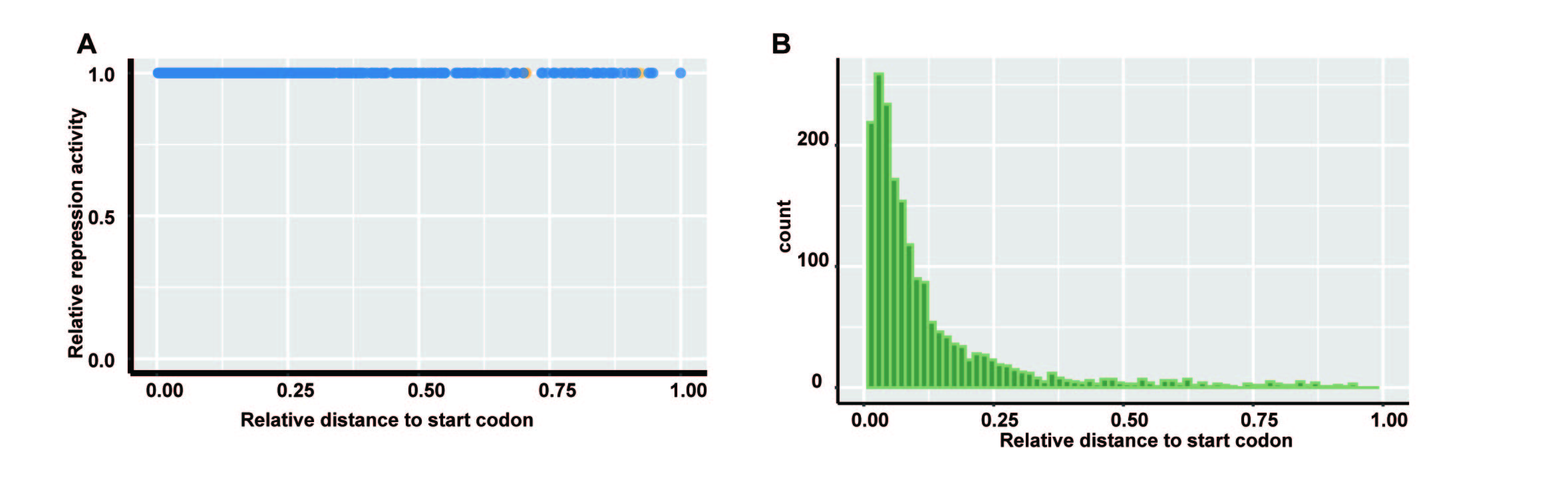

### Figure S3

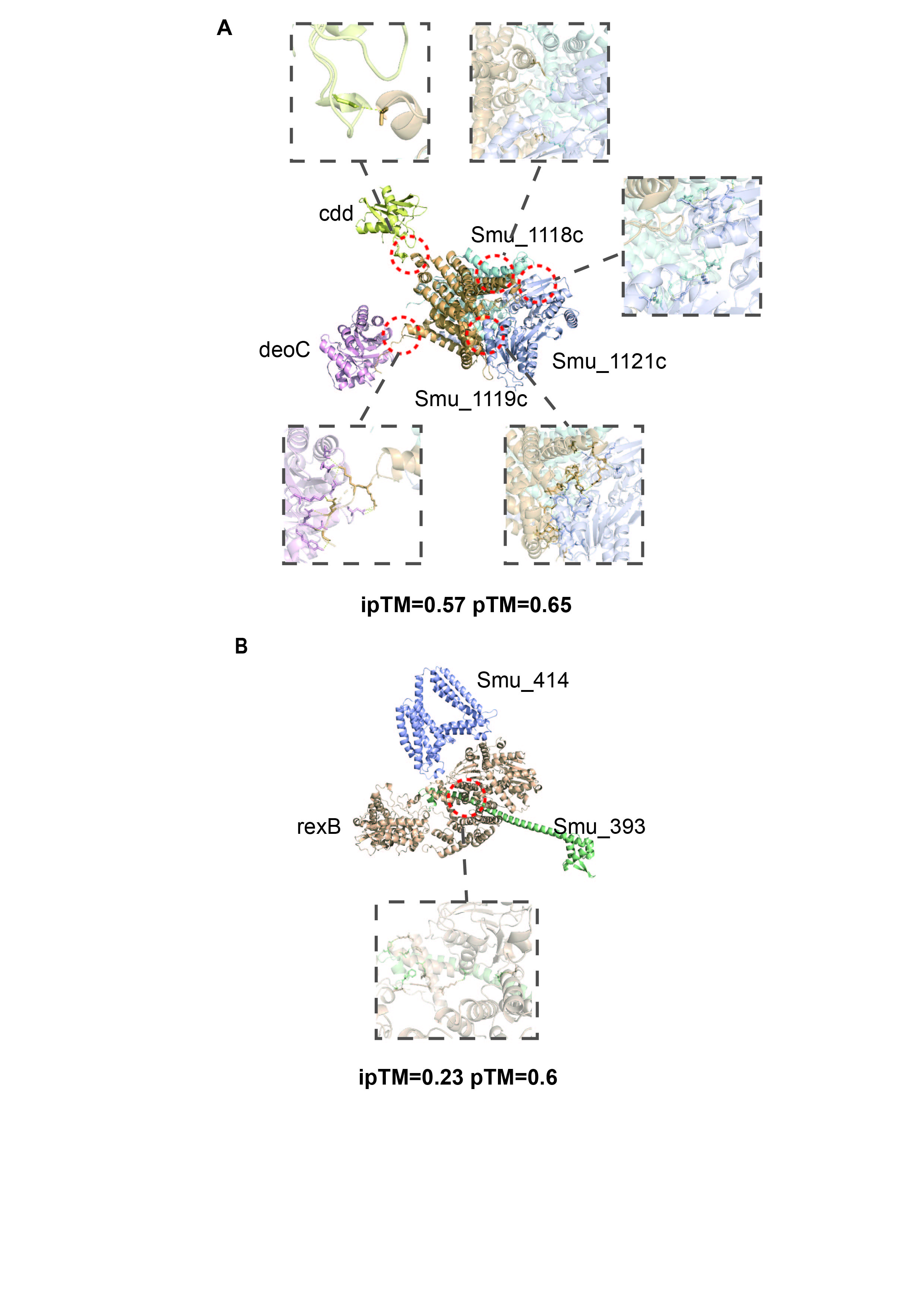
